# GTP-dependent formation of straight oligomers leads to nucleation of microtubules

**DOI:** 10.1101/2020.03.05.979989

**Authors:** Rie Ayukawa, Seigo Iwata, Hiroshi Imai, Shinji Kamimura, Masahito Hayashi, Kien Xuan Ngo, Itsushi Minoura, Seiichi Uchimura, Tsukasa Makino, Mikako Shirouzu, Hideki Shigematsu, Ken Sekimoto, Benoît Gigant, Etsuko Muto

## Abstract

Microtubule (MT) nucleation is essential for cellular activities, but its mechanism is not known because of the difficulty involved in capturing rare stochastic events in the early stage of polymerization. In cells, MTs are nucleated at tubulin concentrations significantly lower than those required for spontaneous nucleation in vitro. The high efficiency of nucleation is due to the synergistic effects of various cellular factors, but the underlying mechanism has not been clarified yet. Here, combining negative stain electron microscopy and kinetic analysis, we demonstrate that the formation of single-stranded straight oligomers with critical size is essential for nucleation in vitro. While the single-stranded oligomers of GTP-tubulin that form prior to MT nucleation show variable curvatures including a few straight oligomers, only curved oligomers are observed among the GDP-bound counterparts. The Y222F mutation in β-tubulin increases the proportion of straight oligomers and drastically accelerates MT nucleation. Our results support a model in which GTP binding causes a small shift in the distribution of oligomer curvature, generating a minor population of straight oligomers compatible with lateral association and further growth to MTs. Our study suggests that cellular factors involved in nucleation promote it via stabilization of straight oligomers.

---

Dynamic arrangement of MT organisation in proper timing and location is critical for various cellular functions, such as cell shape determination and chromosome segregation. The high degree of plasticity observed during MT organization relies on the nucleation and polymerization/depolymerization of individual MT filaments. While catastrophic transition to depolymerization has been well characterized as “dynamic instability”^1,2^, the inverse process involving the nucleation of MTs from tubulin is poorly understood. Nucleation, in general, is a stochastic process, in which the thermodynamically unfavourable growth of pre-critical molecular composites turns into thermodynamically favourable growth via the formation of critical nuclei^3,4^. Because the nucleation intermediates around the critical nuclei appear only transiently, capturing them is a major challenge.

In cells, MTs are nucleated at tubulin concentrations significantly lower than those required for nucleation in vitro^5^. The efficiency of nucleation is increased by the template of the γ-tubulin ring complex (γTuRC) and other cellular factors^6-15^, but the molecular mechanism allowing highly efficient nucleation is unknown. Despite its high efficiency, the kinetics of MT assembly show a time lag between the onset of the reaction and the start of MT growth from the template^6,7^, suggesting that even in the presence of a template, the initial phase of assembly is thermodynamically unfavourable until the nucleation intermediate reaches a critical size.

In this study, we addressed fundamental questions about spontaneous nucleation in vitro, which are also related to nucleation in vivo:(1) What is the size and shape of the critical nucleus, and what happens after its formation? (2) How does GTP binding enable tubulin to nucleate MTs? Because GTP-tubulin assumes a curved conformation in solution^16,17^ and a straight conformation when integrated in the MT lattice^18,19^, during some step in polymerization, GTP-tubulin is expected to change its conformation. However, when and how straightening occurs has been a subject of long debate^17,20,21^. To address the above questions, we combined multiple approaches, including atomic structural analyses of tubulin, statistical analyses of the oligomer shape and macroscopic kinetic analyses.

Our analyses showed that the GTP-dependent formation of straight oligomers with a critical size is essential for the nucleation of MTs. Both GTP- and GDP-tubulin assembled into single-stranded oligomers of various lengths and curvatures, but it was only for GTP tubulin that a minor population of straight oligomers appeared. The proportion of straight oligomers was increased by Y222F mutation in β-tubulin, which also led to an acceleration of nucleation. These straight oligomers of a critical size are compatible for lateral association and further growth to MTs. In vivo, various cellular factors may increase the nucleation efficiency by causing a small change in the proportion of straight oligomers.

## Y222F mutation accelerates nucleation

It has been proposed that GTP binding promotes MT assembly by fostering a conformational switch of the β-tubulin T5 loop, a loop involved in tubulin-tubulin longitudinal contacts (Fig. 1a,b)^22^. The crystal structure of GDP-tubulin indicated that the T5 loop fluctuates between two conformations (“in” and “out”; Fig. 1a). Upon GTP binding, the interaction of the residue D177 in T5 loop with the residue Y222 from H7 helix is broken, and thus the “out” conformation is favoured. By introducing the Y222F mutation, we aimed to produce a stronger bias towards the “out” conformation and to examine how this modulation affects nucleation. To this end, we took advantage of the recent development of a method for the expression and purification of recombinant human tubulin^23,24^. Here, we modified our original method for the preparation of human α1β3 tubulin to purify *Drosophila* α1β1 tubulin. *Drosophila* tubulin has several technical advantages over human tubulin: its yield from host insect cells is better, and it can polymerize at room temperature. Both *Drosophila* WT and Y222F mutant tubulins were produced and purified (Extended Data Fig. 1).

**Fig. 1.**
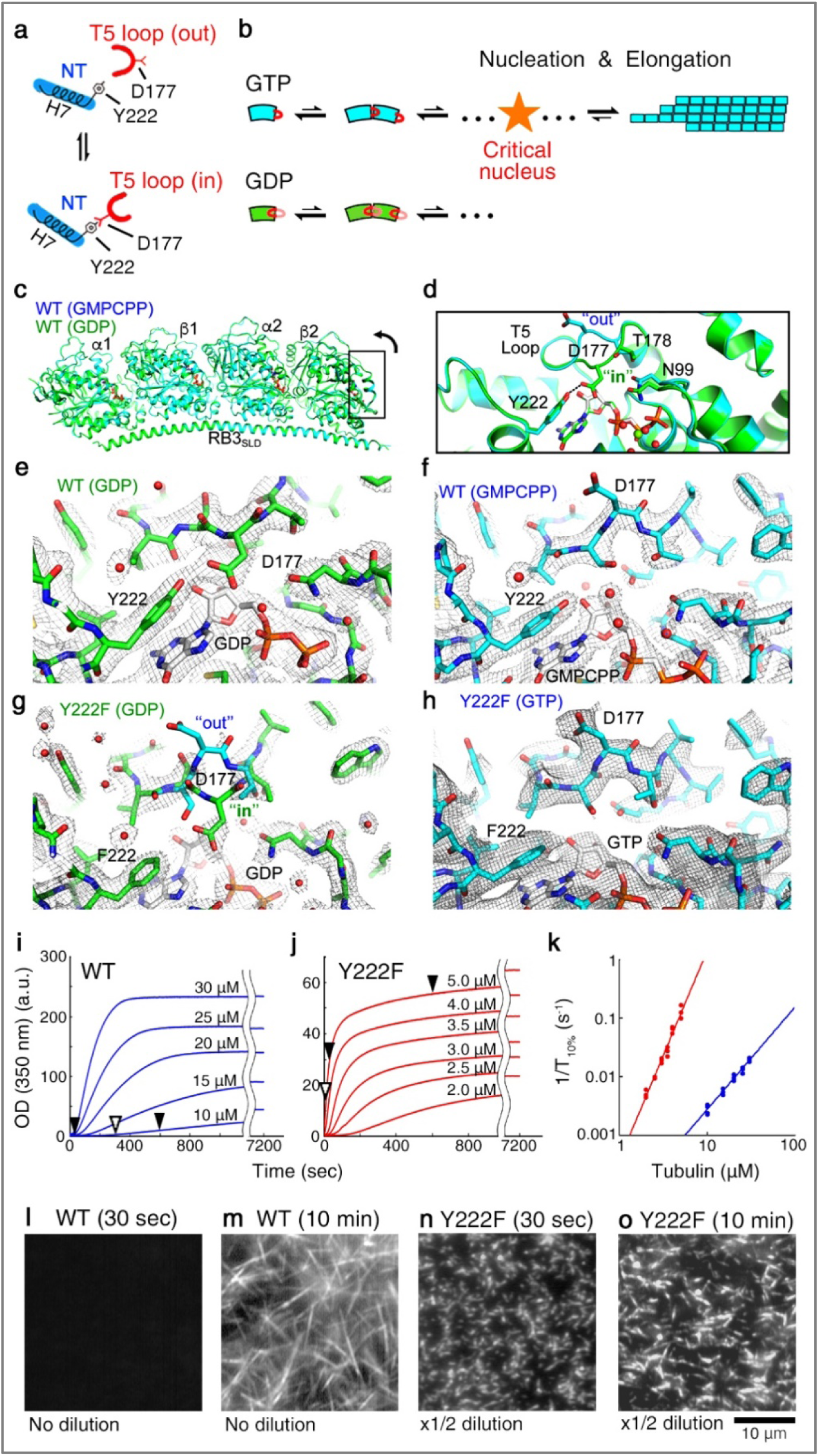
The Y222F mutation modulates the structure of the T5 loop and accelerates nucleation. **a.** Schematic picture of the nucleotide-dependent regulation of the T5 loop in β-tubulin^22^. The “in”/”out” conformation of the T5 loop is regulated by formation/dissociation, respectively, of a hydrogen bond between the D177 residue in the T5 loop and the Y222 residue in H7. GTP/GDP binding biases the structural equilibrium of the T5 loop towards the “out”/”in” conformation, respectively. NT: nucleotide. **b.** Schematic picture of the nucleotide-dependent regulation of the pathways of oligomer assembly. Only for GTP-tubulin with the T5 loop in the “out” conformation does growth become thermodynamically favourable, as the oligomer exceeds the critical size. **c–h.** *Drosophila* WT (**c–f**) and Y222F (**g, h**) tubulin structures within the T_2_R complex. (**c, d**). Superposition of the GMPCPP- and GDP-tubulin structures (**c**, overview; **d**, close-up of the part framed in panel **c** rotated 90 degrees). (**e–h**) 2 F_obs_-F_calc_ electron density maps contoured at the 1 sigma level and centred on the β2 nucleotide binding site and T5 loop. In the case of Y222F(GDP) tubulin, the T5 loop residue D177 and neighbouring residues are not defined in the electron density map, suggesting that this loop is mobile. Both the “in” and “out” conformations of the T5 loop are drawn as a reference. **i–j.** Time course of the polymerization of WT (**i**) and Y222F tubulin (**j**) monitored by OD_350_. At times indicated by black arrowheads, the MTs were checked under the darkfield microscope (panel **l–o**) and negative stain EM (Fig. **2a–c**). At times indicated by white arrowheads, the oligomers were imaged using negative stain EM (Fig. **2e–i**, Fig. **3d–h**). **k.** Log-log plot of the inverse of the time required for the turbidity to reach 10% of its plateau value (1/T_10%_) vs initial tubulin concentration^25^. **l–o.** Darkfield microscopic images of WT (**l, m**) and Y222F MTs (**n, o**). In panel **m**, the WT MTs in solution are shown, whereas in panels **n** and **o**, the Y222F MTs attached on the glass surface are shown. Because of the high MT densities, we could not record the images of the Y222F MTs in solution using darkfield microscopy.

The crystal structure of *Drosophila* WT tubulins within the T_2_R complex showed the nucleotide-dependent movement of the T5 loop in the β2 subunit; the loop was in the “out” and the “in” conformations in the GTP-like and GDP states, respectively (Fig. 1c–f, Extended Data Table 1). This is similar to the nucleotide-dependent structural change of the T5 loop observed for mammalian brain tubulin^22^, though the structural difference between the two states is more pronounced in *Drosophila* tubulin. In the Y222F mutant, whereas the T5 loop was “out” when bound to GTP (Fig. 1h), the loop was poorly defined in the electron density maps in the GDP state (Fig. 1g). Therefore, the Y222F mutation destabilized the T5 loop “in” conformation, but in the absence of GTP, it did not force this loop into the “out” conformation.

We next compared the time course of the assembly of WT and Y222F tubulin by monitoring the turbidity of the tubulin solution (Fig. 1i, j). Because Y222F GTP-tubulin readily polymerized as soon as we transferred the tubulin solution from 4°C to 25°C, we had to monitor the assembly at concentrations significantly lower than the concentrations used for WT tubulin (2–5 μM and 10–30 μM for Y222F and WT tubulin, respectively). Despite the lower concentration range employed, Y222F tubulin assembled much faster than WT tubulin. For both proteins, the turbidity plateau was a linear function of the initial tubulin concentration, yielding critical concentrations (x-intercept) of 4.7 ± 0.5 and 0.19 ± 0.15 μM (mean ± s.d.) for WT and Y222F tubulin, respectively (Extended Data Fig. 1d).

The rate of nucleation was assessed from the inverse of the time required for the turbidity to reach 10% of the plateau value (1/T_10%_)^25^, which was log-log plotted as a function of the initial tubulin concentration (Fig. 1k). For Y222F tubulin, 1/T_10%_ increased in the lower concentration range and showed higher sensitivity to the tubulin concentration compared to the WT, suggesting that Y222F tubulin nucleates faster than WT tubulin^26^. The acceleration of nucleation was also confirmed by using darkfield microscopy. Immediately after the onset of the reaction (at 30 sec), while the Y222F mutant produced many short MTs (initial tubulin concentration: 5 µM; Fig. 1n), no MTs were observed for the WT (10 µM; Fig. 1l). At 10 min, WT MTs were observed but they were fewer in number and longer than the Y222F MTs (Fig. 1m, o). Because of the very rapid nucleation of the Y222F mutant, 5 µM Y222F tubulin and 10 µM WT tubulin were the closest concentrations we could compare.

## Y222F mutation favours straight oligomers

In the above pair of experiments (5 µM Y222F and 10 µM WT), we next analysed the oligomers that coexisted with the MTs by using negative stain electron microscopy (EM). While EM observation requires the sample to be at a low concentration on the order of sub-μM or less^27^, the use of the rapid flush method allowed us to quickly dilute the sample (< 30 ms)^28,29^ and to capture images of the oligomers in the solution used for turbidimetry (Fig. 2a–c, Extended Data Fig. 2). For both WT and Y222F tubulin, oligomers existed at the early stage of assembly when the turbidity was rising, but their numbers declined when the turbidity reached a plateau, suggesting a possibility that these oligomers might have included on-pathway intermediates crucial for MT nucleation. The comparison between the GTP- and GDP-oligomers should provide information on the structural pathway of nucleation (Fig. 1b).

**Fig. 2.**
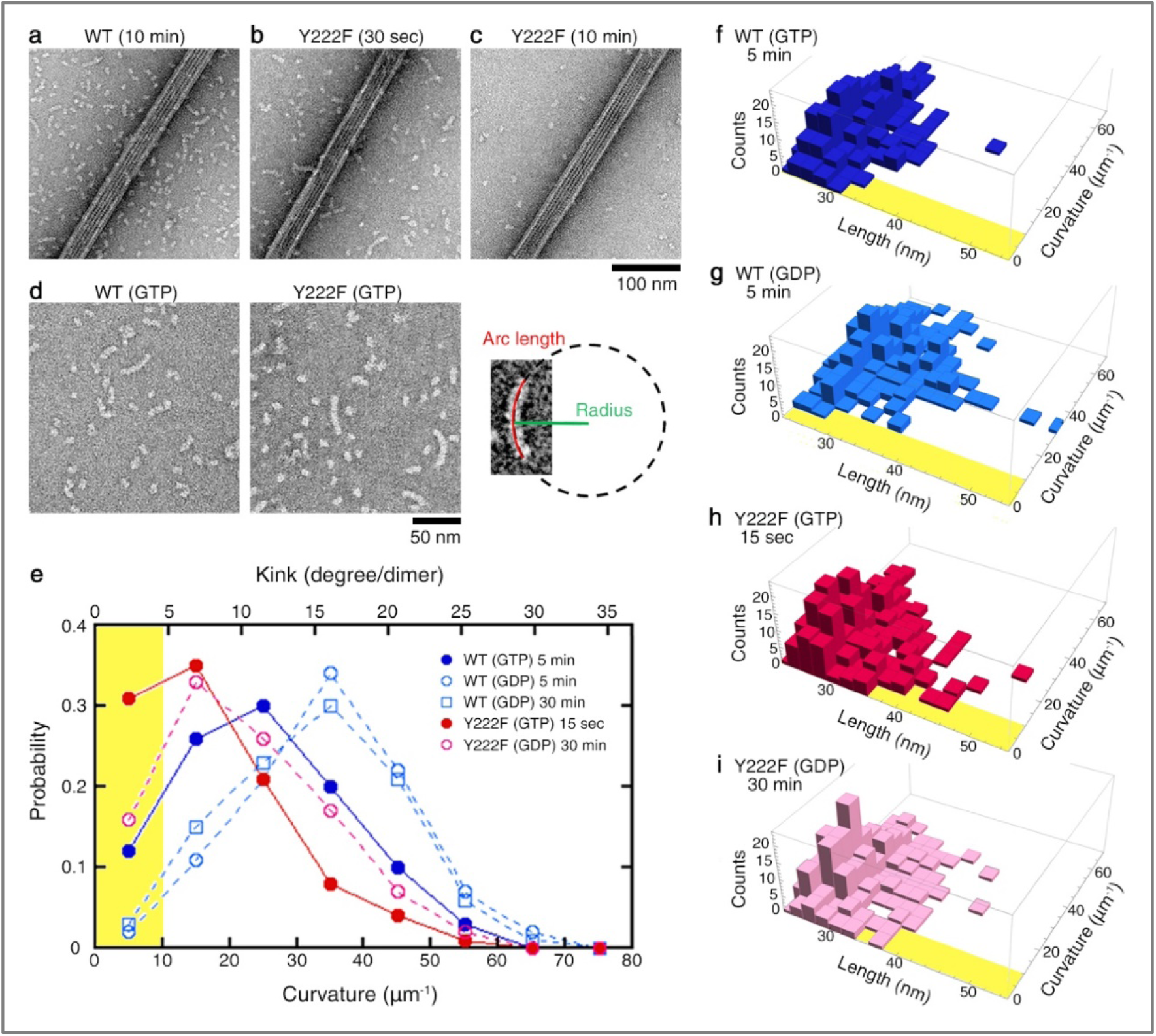
Analyses of oligomer curvature and length by negative stain EM. **a–c.** Electron micrographs of oligomers coexisting with MTs. At 10 min after the onset of reaction, the EM image of the WT showed dimers and oligomers coexisted with the MTs (**a**). For the Y222F mutant, oligomers were observed at 30 sec (**b**) but not at 10 min (**c**). For the darkfield microscopic images of the same samples, see Fig. 1 m–o. **d.** Oligomer length and average curvature were determined by fitting a circle to the centre line of the oligomer. To unambiguously determine the curvature, only those oligomers comprising at least 3 heterodimers were subjected to the analyses. **e.** The normalized distributions of the curvatures. **f–i.** 2D histograms showing the distribution of the length and curvature for WT(GTP), WT(GDP), Y222F(GTP) and Y222F(GDP) oligomers (N = 323, 321, 337 and 300, respectively). In panel **e–i**, the area corresponding to nearly straight oligomers with curvature < 10 µm^-1^ is highlighted in yellow.

As we measured the length and curvature of the WT and Y222F oligomers that appeared in the early stage of assembly (Fig. 2e–i; sampling time 5 min and 15 sec, indicated by the white arrowheads in Fig. 1i and j, respectively), both the WT and Y222F oligomers showed a broad range of curvatures, with the Y222F oligomers being less curved than the WT oligomers (mean ± s.d.: 25.0 ±12.4 and 16.9 ± 10.8 μm^-1^ for WT and Y222F, respectively; Mann-Whitney U test, p<0.01). We also recorded the curvature distributions of oligomers assembled in the GDP condition^30,31^. The GDP oligomers were found to be more curved than the GTP oligomers for both WT and Y222F (mean ± s.d.: 34.4 ± 12.2 and 22.0 ± 11.8 μm^-1^ for WT and Y222F, respectively; Mann-Whitney U test, p<0.01 in both WT and Y222F pairs). The distributions of the curvatures appeared similar to what was reported for the protofilaments at the growing ends of MTs^32,33^.

It is particularly important that a subpopulation of nearly straight oligomers was observed in the presence of GTP (with a curvature < 10 μm^-1^, highlighted in yellow in Fig. 2e). The appearance of these near-straight oligomers correlates with the nucleation rate. With Y222F(GTP) tubulin showing rapid nucleation (Fig. 1j), 31% of the oligomers were nearly straight, whereas with WT(GTP) tubulin showing moderate nucleation (Fig. 1i), only 12% of the oligomers were nearly straight. For WT(GDP) tubulin incapable of nucleation, such oligomers comprised less than 2% of the total.

Notably, 16% of the Y222F(GDP) oligomers were nearly straight, which is comparable to the proportion observed for the WT(GTP) oligomers. Given that the T5 loop in Y222F(GDP) was not anchored in the “in” conformation (Fig. 1g), it implies a possibility that Y222F(GDP) tubulin nucleates MTs. Indeed, incubating Y222F(GDP) tubulin in the condition of the turbidimetry assay led to a signal that slowly increased over time (Extended Data Fig. 4). At 2 h after the onset of the reaction, using darkfield microscopy, we observed the Y222F(GDP) MTs, which were substantially longer and fewer in number than the Y222F(GTP) MTs. Although GTP is required for canonical nucleation under natural conditions, the straight oligomers artificially synthesized by mutation partially complemented the lack of gamma phosphate (γ-Pi).

## Single-stranded oligomers are subcritical

The correlation between the proportion of straight oligomers and the nucleation rate indicates that straight oligomers are essential components of MT nucleation. How do these straight oligomers relate to the critical nucleus? Do they exceed the critical size to enter a stage where growth is thermodynamically favourable? To answer these questions, we calculated the size of the critical nucleus from the turbidity curves and found that the majority of the oligomers we analysed in Fig. 2e were smaller than the critical size, as explained below.

We related the tubulin concentration dependence of polymerization to the size of the critical nucleus. The turbidity (Fig. 1i, j) was first converted to the amount of tubulin in the MTs (Fig. 3a, b), taking the turbidity coefficient into account (see Methods for details). The theory of nucleation predicts that this polymer mass should increase quadratically with time in the early stage of polymerization because both the number of MTs and the masses of individual MTs after nucleation increase linearly with time (Eq. 2 in Methods). Our turbidity data fit well to the quadratic profile, supporting this view. The coefficient of the quadratic function should then be half of the nucleation rate, *I*_0_, multiplied by the MT growth rate, *v*_0_. Substituting *v*_0_ with the actual growth rate (Extended Data Fig. 5), we found that *I*_0_ increases with the α-th power of the initial tubulin concentration *C*_0_, with α being 4.0 ± 0.2 and 5.9 ± 0.3 tubulin dimers (mean ± errors of fit) for WT and Y222F tubulin, respectively (Fig. 3c). α can be interpreted as the minimum size of the oligomers destined to grow, i.e., the size of the critical nucleus. Below this size, the oligomers are in quasi-equilibrium and are undergoing stochastic growth and degradation. As we compared these sizes with the actual length of the oligomers (Extended Data Fig. 6a–f), for both WT and Y222F tubulins, a very large majority of the oligomers were found to be below the critical size. Our results collectively imply that it is the *straight* oligomers of the size α that are critical nuclei for MT assembly.

**Fig. 3.**
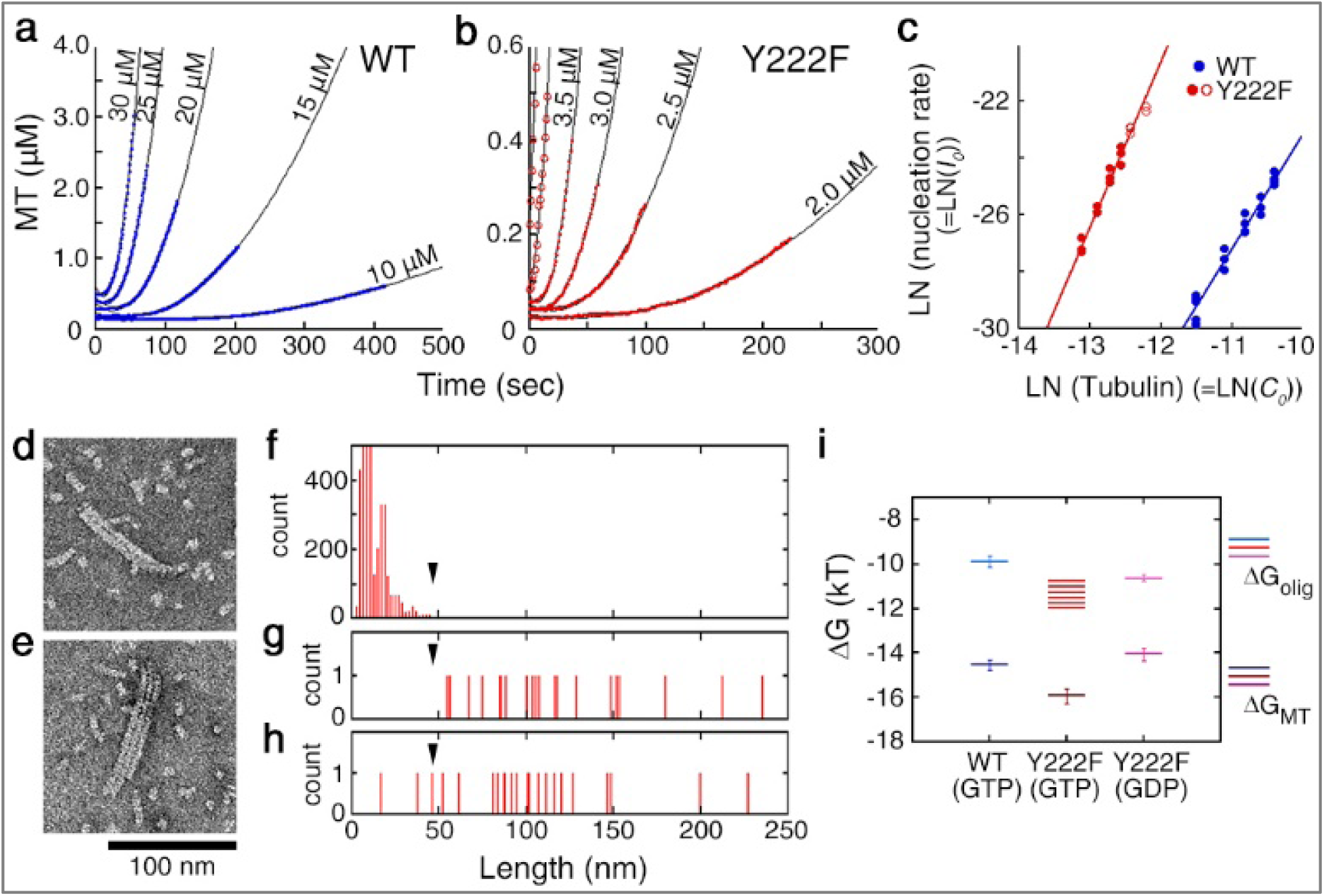
Oligomers below and above the size of the critical nucleus. **a–b.** Time course of the polymerization of WT (**a**) and Y222F tubulin (**b**) from time zero to T_10%_, which was fit by the quadratic function of time (R>0.99 for all curves). The original data are shown in Fig. **1i** and **j. c.** Log-log plots of the rate of nucleation, *I*_0_, vs the initial tubulin concentration, C_0_ (see Methods). For the Y222F mutant, the data measured at 4 and 5 μM tubulin (**○**) were excluded from the calculation of α (see Methods for details). **d–e.** Negative stain electron micrographs of double- or triple-stranded Y222F(GTP) oligomers observed at 15 sec after the onset of reaction. **f.** Length distribution of the single-stranded oligomers (N = 12, 666), which is identical to the dataset shown in Extended Data Fig. **6b, e**. **g–h.** Length distribution of the longer (g) and shorter protofilaments (h) in the two-stranded oligomers (N = 22). The arrowheads in panel **f–h** indicate the size of the critical nucleus (∼6). **i.** The free energy change associated with the binding of the tubulin dimer to the oligomer (ΔG_olig_) and to the MT (ΔG_MT_). For the calculation of errors, see Methods.

From the length distribution of the subcritical oligomers (Extended Data Fig. 6j–l), we also calculated the free energy change associated with tubulin binding to an oligomer, ΔG_olig_ (see Methods for the calculation). The results showed that the interdimer interaction was stronger in the Y222F oligomer than that in the WT oligomer by 1-2 *k*_B_*T* (Fig. 3i; ΔG_olig_ = – 9.9 ± 0.2 *k*_B_*T* and in the range of –10.8 to –12.0 *k*_B_*T* for WT(GTP) and Y222F(GTP) oligomers, respectively). In the Y222F mutant, the interdimer interaction might have been strengthened by the T5 loop being in the “out” conformation^22,34^.

## Oligomers above the critical size

In the above experiment with Y222F(GTP) tubulin, we rarely observed double- or triple-stranded oligomers among the vast majority of single-stranded oligomers (Fig. 3d, e, Extended Data Fig. 7). In the double-stranded oligomer, the length of the longer strand was > 50 nm (Fig. 3g), clearly exceeding the size of the critical nucleus, and also the maximum size of single-stranded oligomers (Fig. 3f). The shorter strands had lengths both above and below 50 nm (Fig. 3h). These results indicate that a single-stranded oligomer that reaches the size of the critical nucleus may laterally associate with a dimer or another oligomer, and continue to grow as a multistranded oligomer. In a later stage (30 sec or longer), we observed sheets composed of a few protofilaments that were several hundred nm in length, likely on their way to becoming MTs (Extended Data Fig. 7)^15,25^.

We could not find multistranded oligomers of WT tubulin (10 μM at 5 min). It is not surprising that we could not find such WT oligomers, considering that the nucleation rate of WT tubulin was approximately three orders of magnitude smaller than that of Y222F tubulin (the nucleation rates in Fig. 3c were 2.1 × 10^−10^ and 2.1 × 10^−13^ M/s for 5 μM Y222F and 10 μM WT tubulin, respectively). Under conditions where nucleation does not occur, for example, in tubulin solution incubated at 4°C (Extended Data Fig. 3), we never observed double- or triple-stranded oligomers.

## Y222F mutation suppresses catastrophe

Y222F tubulin favouring a straight conformation is more efficient than WT tubulin not only in nucleation but also in elongation of MTs. As we compared the dynamic instability of WT and Y222F MT in the GTP condition, the Y222F mutant showed a reduced catastrophe frequency and higher growth rate (Extended Data Fig. 5). Y222F(GDP) tubulin capable of nucleation (Extended Data Fig. 4) was also capable of elongation (Extended Data Fig. 5d, e), although at a rate lower than that of Y222F(GTP) or WT(GTP) tubulin.

From the concentration dependence of the growth rate, we calculated the free energy change associated with the binding of tubulin to the end of an MT, ΔG_MT_ (Fig. 3i, see Methods for calculation). In both WT(GTP) and Y222F(GTP), tubulin was ∼4 *k*_B_*T* more stabilized upon integration into the MT lattice than upon binding to the end of a single-stranded oligomer (ΔG_olig_). This difference is likely attributed to the energy gain due to the lateral interaction, as was reported for brain tubulin^35,36^.

## Mechanism of GTP-dependent nucleation

By linking the statistical analysis of pre- and post-critical nuclei with the kinetic analysis of the MT assembly, we propose the following scheme for GTP-dependent nucleation of MTs. The dimers and sub-critical oligomers of variable sizes and curvatures are in quasi-equilibrium in tubulin solution (Fig. 4a). The GTP-dependent extension of the T5 loop stabilizes the longitudinal interdimer interaction^22,34^, giving rise to a subpopulation of nearly straight oligomers (highlighted in yellow in Fig. 2e–i). The oligomers also laterally interact and transiently form double-stranded oligomers, but most of them dissociate almost immediately because of the weakness of the lateral interaction compared to the longitudinal interaction^18,37,38^. With increasing oligomer size, the population decays exponentially (Extended Data Fig. 6j), whereas the potential to form double-stranded oligomers increases owing to the increased lateral interface. When a single-stranded oligomer reaching sufficient size and straightness (= critical nucleus) laterally associates with a dimer or another oligomer, the consequential two-stranded oligomer will serve as a platform for the growth of protofilaments (= seed; Fig. 4b, lower panel).

**Fig. 4.**
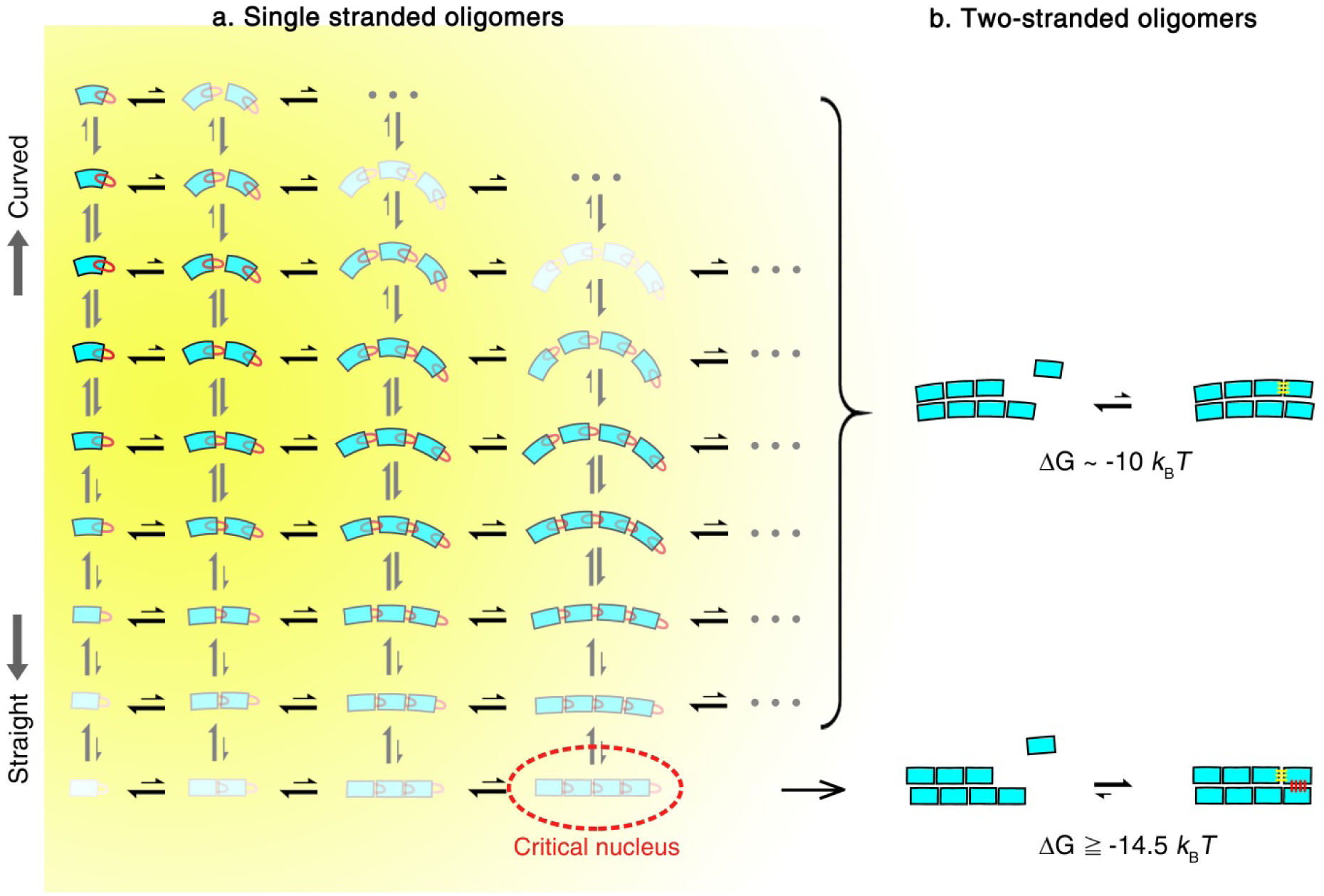
A model for the structural pathway of spontaneous nucleation. **a.** In a solution of GTP tubulin, tubulin dimers and oligomers of variable sizes and curvatures are in rapid equilibrium. The colour intensity represents the probability for each dimer and oligomer. The distance between tubulin dimers in an oligomer symbolically represents the strength of the interdimer bonds. Straight oligomers crucial for nucleation can be formed only from GTP-tubulin, with the T5 loop “out” conformation being favoured. **b.** While the reaction of tubulin binding to a two-stranded oligomer composed of straight protofilaments can be thermodynamically favourable (lower panel), the reaction of tubulin binding to a two-stranded oligomer formed from curved protofilaments is not (upper panel). Here, we assume that a tubulin dimer that binds to a multistranded oligomer gains energy equivalent to the energy gained by a tubulin dimer that binds to an MT protofilament (ΔG_MT_).

The binding of a tubulin molecule to such a multistranded oligomer might be thermodynamically more favourable than binding to single-stranded oligomers, because in the former case, simultaneous longitudinal and lateral interactions are possible. Our analyses showed that the free energy change associated with the tubulin binding to MT is 4 *k*_B_*T* larger than the free energy change associated with the binding to a single-stranded oligomer (Fig. 3i). Similarly, tubulin that binds to multistranded oligomer would gain an energy equivalent to this 4 *k*_B_*T* (Fig. 4b, lower panel). For a multistranded oligomer composed of curved oligomers, if any, such a thermodynamic transition is difficult because an incoming tubulin would participate only in the longitudinal interaction (Fig. 4b, upper panel).

According to this scheme, nucleation depends on a very rare event, i.e., the formation of a straight oligomer with a critical size. That both GTP- and GDP-tubulin assumed a similarly curved conformation in the conditions the assembly did not occur^16,17,22,39^ is not contradictory to our data because straight oligomers are a very minor population that appear only in the MT assembly condition.

The Y222F mutation bringing a bias in the structural equilibrium of oligomers towards a straight conformation caused rapid nucleation and the reduction of the catastrophe frequency. This behaviour parallels the behaviours of MTs observed in the presence of GMPCPP and Taxol^40-44^. Both ligands are known to shift the distribution of the oligomer/protofilament curvature towards a straight conformation^40,45^. It is possible that any factor that stabilizes straight oligomers/protofilaments accelerates nucleation and prevents catastrophe.

The features of MT nucleation in cells are reminiscent of those described here for nucleation in vitro. In particular, the power law dependence of the γTuRC-mediated nucleation on tubulin concentration (Ref. 14 and Extended Data Fig. 8) strongly suggests that templated nucleation also requires tubulin to assemble to a critical size, as was shown for the spontaneous nucleation (this study and Ref. 25). In addition, nucleation is enhanced by MT-associated proteins (MAPs) that promote MT growth (e.g., XMAP215)^6,7,11,12^ or suppress catastrophes (e.g., TPX2)^6-8^, as the Y222F mutation does. Whether the MAPs do so by increasing the population of straight oligomers remains to be determined. In testing this hypothesis, the methods of analysis developed in this study will become a powerful tool. The past models of dynamic instability^2,20,46^ and of nucleation have scarcely considered the diversity of conformations of tubulin and oligomers. Our study offers a new viewpoint focusing on the rare events.

## Methods

### Construction of baculovirus transfer plasmids

*Drosophila* α1-tubulin (αTub84B; NT_033777) and β1-tubulin (βTub56D; NT_033778) genes were obtained from the Drosophila Genomics Resource Centre. For affinity purification, tags were fused to the 3’ end of each clone (for α1-tubulin, a His_8_ tag and a sequence encoding FactorXa cleavage site (IEGR) were linked by a glycine-based linker sequence (GGSGG), and for β1-tubulin, a FLAG tag and an IEGR were linked by a linker sequence (GGG)). To increase the expression level, an L21 leader sequence was also added just before the start codon^47^. To exclude variability due to acetylation, the α1 was made unacetylatable by residue substitution K40R, and it was treated as wild type. The inserts were cloned into the pFastBac Dual vector (Life Technologies)^23^.

### Purification of recombinant tubulin

The recombinant tubulin was expressed in HighFive cells (Life Technologies) and purified by three steps of column chromatography (DEAE Sepharose ion exchange chromatography, His-affinity column, FLAG-affinity column), as previously described^23^, with the exception of the following: While 1 mM GTP was included in solutions used in all three steps of the purification of WT tubulin, GTP was replaced by 1 mM GDP for purification of Y222F tubulin (Extended Data Fig. 1). The tubulin eluted from the FLAG-affinity column was concentrated to > 5 mg/mL with an Amicon Ultracel-30K filter (Millipore) centrifuged at 100,000 rpm (∼540,000 xg, Beckman TLA-100.3 rotor) for 10 min at 4°C, and the supernatant was incubated at 30°C after adding Glycerol (final 33% v/v) and GTP (2 mM). After 2 h of incubation, Factor Xa protease (New England Biolabs) was added to the MT solution at a molar ratio of 1 : 22 (Factor Xa : tubulin) and reacted overnight at 25°C. The next morning, the sample solution was centrifuged at 100,000 rpm (Beckman TLA-100.3 rotor) for 15 min at 30°C, the precipitated MTs were suspended in BRB80 buffer solution (80 mM PIPES, 2 mM MgCl_2_, 1 mM EGTA) containing 1 mM GTP (for WT tubulin), or 1 mM GDP (for Y222F tubulin) at 4°C, and left for 30 min on ice. To remove aggregates, the solution was centrifuged again at 100,000 rpm (∼430,000 xg, Beckman TLA-100 rotor) for 10 min at 4°C. The supernatant was frozen in aliquots in liquid nitrogen and they were stored at –80°C until use. The yield of tag-free tubulin was ∼3 mg from a 1-L culture of HighFive cells (∼20 g of cells). Protein concentration was determined using Pierce^TM^ 660nm Protein Assay Kit (Thermo Fisher Scientific).

### Preparation of GTP-tubulin for turbidimetry and negative stain EM

To follow the pre-equilibrium dynamics upon nucleation of MTs (Figs. 1–3), rigorous control of the starting material was important. To make sure that at time zero the oligomers are in quasi-equilibrium with the dimers at 4°C, the tubulin sample was prepared fresh for each run of the experiment by the following protocol: Soon after the frozen tubulin sample was defrosted, it was filtered by an Ultrafree-MC VV Centrifugal filter (0.1 μm pore size; Millipore) to remove aggregates, and then spun through a Micro Bio-Spin P30 Column (Bio-Rad Laboratories) to replace the solution by BRB80 buffer containing 1 mM GTP. A 70-μL tubulin fraction eluted from the column was centrifuged at 100,000 rpm (Beckman TLA-100 rotor) for 15 min at 4°C to remove aggregates and diluted at the appropriate concentration. The tubulin sample was used for turbidimetry and negative stain EM within 30 min after the ultracentrifugation (Extended Data Fig. 3). The nucleotide content in the E-site was > 95% GTP (Extended Data Fig. 1b).

### Preparation of GDP-tubulin for turbidimetry and negative stain EM

Soon after being defrosted, tubulin was first filtered by an Ultrafree-MC VV Centrifugal filter (0.1 μm pore size; Millipore) to remove aggregates. For Y222F tubulin, the sample was further spun through a Micro Bio-Spin P30 Column (Bio-Rad Laboratories) to replace GDP with GTP. WT and Y222F tubulins, thus conditioned, were incubated at 30°C for 2 h for polymerization. The sample solution was centrifuged on a 60% glycerol cushion at 100,000 rpm (Beckman TLA-100 rotor) for 15 min at 30°C to precipitate the MTs. After careful washing of the wall of the centrifuge tube and the pellet surface by nucleotide-free BRB80 buffer, the pellet was suspended in nucleotide-free BRB80 buffer at 4°C. The sample was left on ice for 30 min, and a 70-μL sample solution was centrifuged at 100,000 rpm (Beckman TLA-100 rotor) for 15 min at 4°C. The supernatant diluted at the appropriate concentration was used for turbidimetry and for preparation of the grid for negative stain EM. The nucleotide content in the E-site is ∼100% GDP for both WT and Y222F tubulin (Extended Data Fig. 1b). Similar to the preparation of GTP tubulin, the sample was prepared fresh for each run of the experiment.

### Nucleotide content analysis

The tubulin sample was spun twice through a Micro Bio-Spin P30 Column, equilibrated with nucleotide-free BRB80, and was denatured with trifluoroacetic acid (final 2%). After removal of the denatured protein by centrifugation, the supernatant containing the released nucleotide was filtered through an Ultrafree-MC GV Centrifugal Filter (0.22 μm pore size; Millipore) to remove small debris. The filtrate was analysed by fast protein liquid chromatography (AKTApurifier, GE Healthcare Life Sciences) using a MonoQ column (5/50 GL, GE Healthcare Life Sciences) equilibrated with 20 mM Tris-HCl pH 8.0. The nucleotide was eluted using a 0–300 mM NaCl gradient. GDP, GTP, and GMPCPP solutions were injected as references to calibrate the column.

### Cryo-EM sample preparation

The WT and Y222F MTs were assembled at 15 μM and 4 μM of the WT and Y222F tubulin dimers in BRB80 buffer containing 1 mM GTP at room temperature (25–30°C), respectively. The WT and Y222F MTs were applied to lacey carbon grids (Agar Scientific AGS166-4 400 mesh copper grids) and holey carbon grids (Quantifoil R1.2/1.3 300 mesh copper grids) immediately after application of glow discharge to the grids, and were vitrified by using EM GP (Leica) and Vitrobot (Thermo Fisher Scientific), respectively. The grids of the WT and Y222F MTs were observed at liquid nitrogen temperature by using a JEM 2100F electron microscope (JEOL) operated at 200 kV and a Tecnai Arctica electron microscope (Thermo Fisher Scientific) operated at 200 kV, respectively. The EM images of the WT and Y222F MTs were captured by an UltraScan 4000 CCD camera (GATAN) at nominal magnification of 50,000 with 0.213 nm/pixel and by a Falcon II direct electron detector (Thermo Fisher Scientific) at nominal magnification of 39,000 with 0.285 nm/pixel, respectively. The defocus range was between –2.6 μm to –3.5 μm. The total dose was 0.1–0.4 e^-^/nm^2^.

### Crystallisation and structure determination

Crystals of *Drosophila* tubulin were obtained as T_2_R complexes and after seeding^22^. Such complexes comprise two tubulin heterodimers and one stathmin-like domain of the RB3 protein (RB3_SLD_). Data for WT(GDP) tubulin were collected at the ID23-1 beamline (European Synchrotron Radiation Facility, Grenoble, France), data for WT(GMPCPP) tubulin at the Proxima-1 beamline (SOLEIL Synchrotron, Saint-Aubin, France), and data for both Y222F(GDP) and (GTP) tubulin at Proxima-2 beamline (SOLEIL Synchrotron). They were processed with XDS^48^. The sT_2_R structure (PDB ID 3RYC) was used as a starting point for refinement with BUSTER^49^ with iterative model building in Coot^50^. Data collection and refinement statistics are reported in Extended Data Table 1. The atomic coordinates and structure factors have been deposited in the Protein Data Bank under accession codes 6TIS (WT(GDP) tubulin), 6TIY (WT(GMPCPP) tubulin), 6TIZ (Y222F(GDP) tubulin) and 6TIU (Y222F(GTP) tubulin). Figures of the structural models were generated with PyMOL (www.pymol.org).

### Turbidimetry

Polymerization of MTs (in BRB80 buffer containing 1 mM GTP) was monitored by turbidimetry at 350 nm on a spectrofluorometer (FP-6500; JASCO). A micro quartz cell with an optical path of 3 mm (FMM-100, JASCO) was placed in a cell holder maintained at 25 ± 0.2°C. Polymerization was started by introducing a 50-μL tubulin sample, kept at 0°C, into the cell. The temperature of the sample solution reaches 25°C with a relaxation time of 4 sec, when monitored by fluorescent temperature indicator^51^. In both WT and Y222F, the turbidity returned to the original level when the temperature was lowered to 4°C.

### Darkfield microscopic observation of MTs

At a desired time point in the course of turbidity measurement, an aliquot of MT solution was sampled, diluted in BRB80 buffer solution containing 1 mM GTP for optimal observation of individual MT filaments, and introduced into a flow chamber prepared from a coverslip and a glass slide (No. 1 coverslip and FF-001 glass slide, respectively, from Matsunami Glass.) that were spaced by double-sided scotch tape (dimensions 9 mm x 9 mm x 80 μm). The chamber was sealed with VALAP and the sample was observed under a darkfield microscope (BX50, Olympus) equipped with an objective lens (either Plan lens, x40, NA=0.65, or Plan lens, x20, NA=0.4, Olympus) at 25 ± 1°C. To avoid any influence of the glass on nucleation and polymerization^8^, the MTs freely floating in solution were observed. The images were projected onto an image intensified CCD camera (C7190, Hamamatsu Photonics) and stored in a digital video recorder (GV-HD700, Sony). The images were recorded between 30 sec and 2 min after the dilution of MTs, during which timeframes the number and length distribution of the MTs were virtually unchanged. To avoid aggregation of MTs, we did not use the cross-linking reagents to fix MTs, except for the case of the Y222F MTs shown in Fig. 1n, o. In Fig. 1n, o, to resolve the transient short MTs, the MTs were fixed with 1% glutaraldehyde and the images of the MTs attached to the surface of the glass slide were recorded.

### Rapid flush negative stain EM

For WT (either GTP or GDP loaded) and Y222F(GDP), the reaction of polymerization was started by transferring the tubulin sample from a water bath set at ∼0°C to a water bath at 25°C. One minute before the termination of the reaction, 50 μL of 1% uranyl acetate, 7 μL of air, and 0.5 to 3 μL of 10-μM WT or 5-μM Y222F tubulin in BRB80 buffer were sequentially drawn into a pipet tip attached to a Gilson P200 Pipetman (Gilson) at 25°C. At the time point for the termination of the reaction (the exact numbers are indicated in the graph legend for Fig. 2e), the entire contents of the tip were ejected onto a carbon film of an EM grid (Okenshoji). The rapid flush method allows immediate dilution and fixation of the protein samples within 30 ms^28,29^. In the case of Y222F(GTP), the protocol was arranged to keep up with the rapid nucleation. The reaction of polymerization was started by drawing 1–2 μl of 5 μM Y222F(GTP) tubulin (in BRB80 buffer) at 4°C into a pipet tip at 25°C, which was preloaded with 7 μL of air and 50 μL of 1% uranyl acetate solution. Fifteen seconds later, the polymerization reaction was terminated by ejecting the entire contents of the tip onto the EM grid, as described above. In all experiments (WT and Y222F, either GTP- or GDP-loaded), after blotting off the excess stain, the grid was dried at room temperature. The sample was observed with a JEM1400Plus electron microscope (JEOL) operated at 80 kV. The images were recorded on an EM-14800RUBY CCD camera (JEOL) at nominal 40,000 magnification (for the analysis of oligomers; Extended Data Fig. 2e-h) or nominal 2,500 magnification (for examination of MTs; Extended Data Fig. 2a-d), each corresponding to a pixel size of 0.413 or 6.59 nm, respectively.

### Measurement of oligomer curvature and length

For the analyses of curvature (Fig. 2e), dimers and tetramers (i.e., the oligomers with a length below 23 nm) were excluded from the analyses because they are too short for accurate curvature measurement. For each oligomer, the arc length and radius of curvature were manually measured by drawing a polyline along the central trajectory of the oligomers and fitting a circle from the polyline by using ImageJ (https://imagej.nih.gov/ij/; Fig. 2d). From the radius of curvature *r* (nm), the curvature *κ* (μm^-1^) and the kink between the tubulin dimer *θ* (degree) were calculated by using the relationships *κ* = 1000 / *r* and *θ* = 360 * (*l* / 2*nr*), with an average dimer length, *l*, of 8 nm (Fig. 2e–i). The measurement was performed by two independent researchers who were blinded to sample identification.

For quantitation of the length distribution of oligomers (Extended Data Fig. 6a–f), the length of all oligomers in 10–12 micrographs (with size 843 × 847 nm), each covering ∼1,000 oligomers in total, were manually measured using Image J (the exact numbers are indicated in the legend of Extended Data Fig. 6). The experiments were performed twice for each condition (10 μM WT(GTP) at 5 min, 5 μM Y222F(GTP) at 15 sec, and 5 μM Y222F(GDP) at 30 min after the onset of the polymerization reaction). For the oligomers with a length above 12 nm, the central trajectory of the oligomer was fit by a circle and the arc length was measured. For the oligomers with a length below 12 nm, the largest distance between two points within the object (Ferret diameter) was measured.

### In vitro MT dynamics assays

The surface of the glass slides (FF-001, Matsunami Glass) were coated with biotin-functionalized polyethylene glycol (PEG) (BIO-PEG-SC molecular weight = 3,400, Laysan Bio)^52^. A microscope chamber (18 mm x 6 mm x ∼100 μm) was constructed using a coverslip (No. 1 coverslip, Matsunami Glass) and a PEG-coated glass slide. GMPCPP-stabilized and biotinylated MT seeds were prepared from WT tubulin as described in Gell et al.^53^. For dynamic assay, the GMPCPP seeds were immobilized on the bottom of the chamber (= PEG-coated glass) via streptavidin (S888, Thermo Fisher Scientific), and 20 μL of WT or Y222F tubulin in BRB80 buffer solution containing 1 mM GTP was infused into the chamber. The chamber was sealed by vacuum grease and nail enamel, and the sample was observed under a darkfield microscope (BX50, Olympus; Plan lens, x20, NA = 0.4, Olympus) at 25 ± 1°C. Time-lapse images of the MTs were projected onto a high-sensitivity CCD camera (WAT-910HX/RC, Watec) at 1 frame/sec, and the data were stored on a hard disc (DFG/USB2pro converter, The Imaging Source; Latitude 7370, Dell).

### Dynamic parameter measurement

Kymographs were generated from the time-lapse darkfield images of MTs using ImageJ. The MT growth rate was determined from the slope of the growing MT in the kymograph (Extended Data Fig. 5a–e). The mean and standard deviation of growth rate was calculated from all MTs analysed for each condition (the number of MTs analysed per data point was 4 to 44; a total of 217, 122, and 235 MTs were analysed for WT(GTP), Y222F(GTP), and Y222F(GDP) datasets, respectively). From the tubulin concentration dependence of the MT growth rate, *v*_0_[/s] = *k*_+_*C*_0_ − *k*_−_, the rate constants for the association and dissociation of tubulin to the MT plus end (*k*_+_ and *k*_-_) were calculated assuming that a MT with a length of 1 μm corresponds to 1,634 dimers^54^ (Extended Data Fig. 5e, g). Neither WT(GTP) nor Y222F(GTP) tubulin grew onto the minus end, whereas some minus-end elongation was observed with Y222F(GDP) tubulin. The catastrophe frequency was determined as the number of observed catastrophe events divided by the total time spent in the growth phase (363–2491, 112–1596, and 415–576 min per data point for WT(GTP), Y222F(GTP), and Y222F(GDP) tubulin, respectively). The standard deviation was estimated by the catastrophe frequency divided by the square root of the number of catastrophes, assuming the catastrophe events are Poisson processes (Extended Data Fig. 5f, g). Assuming that the catastrophe frequency declines linearly with the tubulin concentration, we estimated the range of tubulin concentration where the catastrophe is suppressed (= x-intercept; Extended Data Fig. 5g).

### Assessment of the size of critical nucleus

Assuming that the turbidity is proportional to the amount of tubulin polymerized into MTs, the turbidity (Fig. 1i, j) was converted to the concentration of tubulin in MTs (Fig. 3a, b) by dividing the OD_350_ values by the turbidity coefficient (= slope of the lines in Extended Data Fig. 1d)^55^, 9.14 and 13.67 for WT and Y222F, respectively. The linearity between the plateau value of turbidity and the initial tubulin concentration justifies this conversion (Extended Data Fig. 1d).

We applied the standard nucleation-and-growth model^26^ to these kinetic curves of nucleation. The oligomers of critical size are, by definition, the minimal assemblies that are destined to grow. As entering into this stage is a very rare event, we supposed that the oligomers not exceeding the critical size are almost in equilibrium (quasi-equilibrium), and are undergoing the stochastic growth and degradation. In that condition, the nucleation rate, *I*_0_ (the rate of increase in the number concentration of critical nucleus), is given by

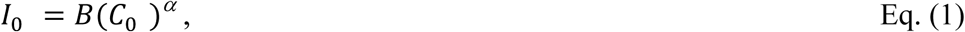

where *C*_0_ is the concentration of the tubulin dimers in solution, α represents the size of the critical nucleus, and *B* is a constant. In the regime where the turbidity is below 10% of the plateau value (*t* < T_10%_), *C*_0_ can be regarded as constant. The meaning of *B* reflects the specifics of the formation and configuration of the critical nucleus. In cases where the critical nucleus is formed by some conformational change of a linear oligomer of size α, *B* would be the equilibrium constant of the assembly of α dimers multiplied by the rate constant of the conformational change. However, in the case where irreversible growth is triggered by the incorporation of the α-th tubulin dimer into the least stable oligomer, *B* would represent the binding rate of the last dimer. In both cases, we have the above form for *I*_0_.

While we know that the single-stranded oligomers and MTs coexist in the solution (Fig. 2a–c, Extended Data Fig. 2), we have little information regarding the reaction cascade between the formation of critical nucleus and the appearance of a tubular form of MT. This passage may take some time, which we tentatively denote by Δ (see below).

Once the tubular form of the MT is achieved, the MT elongates at an average growth rate of *v*_0_, which can be separately measured in MT dynamics assay (Extended Data Fig. 5e). Then the nucleation events taking place in the interval time τ and τ + *dτ* have the concentration *I*_0_*dτ*, and each of those events contributes to the MT mass through the form *v*_0_ × (t – τ-Δ) at time *t* (>τ +Δ). In the early phase of polymerization (*t* < T_10%_), the concentration of tubulin is high enough that we can ignore the catastrophe (Extended Data Fig. 5f, g). Therefore, the total mass of MTs at time *t* is

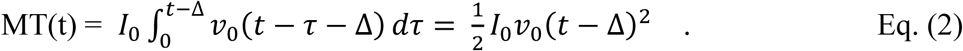

This equation predicts that in the early stage of polymerization, the polymerization curve rises quadratically with time with a quadratic coefficient of 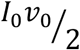. Indeed, polymerization kinetics from time zero to T_10%_ fits very well to the quadratic function (Fig. 3a, b). By substituting the growth rate *v*_0_ measured in a MT dynamic assay (Extended Data Fig. 5), we can find *I*_0_ at each concentration *C*_0_. In Fig. 3c, the log-log plot of *I*_0_ *vs C*_0_ gives a line with a slope corresponding to the size of critical nucleus (α), 4.0 ± 0.2 and 5.9 ± 0.3 for WT (●) and Y222F (●) tubulin, respectively. We checked how the inclusion of the lag time Δ in Eq. (2) affects α. While the introduction of the lag Δ in Eq. (2) improves the quadratic fit (R improved from 0.98 to 0.99 for both WT and Y222F), it affects the value of α by no more than 7%. These results justify our estimation.

In estimating the size of critical nucleus, we were concerned about the possible errors due to the nonlinearity of OD_350_ to the polymer mass of very short MTs. In the above calculation for Y222F tubulin, the data measured for 4- and 5-μM tubulin were not included, because some of the MTs could be too short to assure the linearity between the turbidity and the mass of MTs^56^. When we included the data for 4- and 5-μM tubulin in the fitting (Fig. 3c, ●+○), α was 5.4 ± 0.2, virtually the same as the calculation excluding these two dataset.

The size α calculated by an alternative method also confirms the validity of our method. The assessment of the size from the power law dependence of 1/T_10%_ on tubulin concentration (Fig. 1k)^25^ showed α for WT and Y222F tubulin to be 3.5 ± 0.1 and 6.1 ± 0.4, respectively, close to the values led by our method (4.0 ± 0.2, 5.9 ± 0.3 for WT and Y222F tubulin, respectively). The calculation based on the 10%-criterion (1/T_10%_) slightly underestimates the size α for WT tubulin because at the tubulin concentrations near the critical concentration (for example at 10 μM tubulin), T_10%_ is affected not only by the nucleation and elongation but also by the catastrophe, breaking the premise for this calculation that 1/T_10%_ is determined only by the nucleation and elongation^26^.

The size of the critical nucleus estimated for Y222F tubulin is comparable to the size estimated for brain tubulin where the nucleation was accelerated by Glycerol (α = 7.5), calculated by the power law dependence of 1/T_10%_ on tubulin concentration^25^ (according to their definition of critical nucleus, α = 6.0).

### Assessment of the size of critical nucleus for templated-nucleation

Zheng et al.,(1995)^14^ counted the number of MTs, *N*_MT_, formed in the initial 5 min of incubation of brain tubulin at 37°C both in the presence and absence of γTuRC (Fig. 3 of Zheng et al.). The dependence of this number on the tubulin concentration is fit by the equation

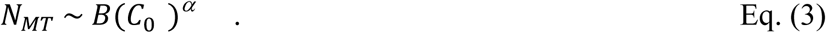

The size of the critical nucleus, α, for the nucleation templated by γTuRC was then estimated as 6.7 ± 0.9 (Extended Data Fig. 8b).

### Calculation of the molar concentration of 2*n*-mer, *x*_2*n*_

Based on the dataset for oligomer length, we first plotted a cumulative frequency of length (Extended Data Fig. 6g–i). These plots were then numerically converted into an empirical probability density as a function of oligomer length. After smoothening the densities using a Gaussian filter, we read off each threshold length separating the first and second, the second and third, and the third and fourth populations, respectively (i.e., 2n-mer and 2(n+1)-mer with n = 1–3). For those thresholds between the oligomers of larger size (n>3), they were determined by extrapolating the average spacing between the thresholds for the smaller sizes (n≤3). The probability of 2n-mer, *p*_2*n*_, was then calculated by integrating the probability density over the interval between neighbouring threshold lengths.

The next step is to connect *p*_2*n*_ to the molecular concentrations (*x*_2*n*_).At the time point when the polymerization reaction was terminated on the EM grids, the turbidity reached 7.0% (WT(GTP)) and 25.3% (Y222F(GTP)) of the plateau value. Considering the initial tubulin concentration (C_0_; 10 and 5 µM for WT(GTP) and Y222F(GTP) tubulin, respectively) and the critical concentration (C_c_; 4.7 and 0.19 µM for WT(GTP) and Y222F(GTP) tubulin, respectively), the concentration of tubulin dimers contained in the ensemble of oligomers 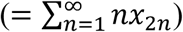 is estimated to be 9.6 and 3.8 μM for WT(GTP) and Y222F(GTP) tubulin, respectively. For Y222F(GDP) tubulin, 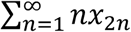 is estimated to be 4.9 μM, taking the polymerized 0.1-μM tubulin (estimated from the turbidity at the time the reaction was terminated) into account. Finally, the molar concentration of each 2n-mer (*x*_2*n*_) was calculated from the total concentration of oligomer, 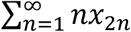, and the probability of each 2n-mer, *p*_2*n*_ (Extended Data Fig. 6j–l). Because the calculation neglects the small population of multistranded complexes or sheets that are too small to contribute the turbidity^56^, the total concentration of oligomers might include some errors, giving only an upper limit for 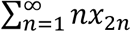.

### Calculation of the specific free energy gains, ΔG_olig_ and ΔG_MT_, upon binding of tubulin dimer

Assuming that the ensemble of pre-nucleus oligomers with various sizes are in rapid quasi-equilibrium, the free energy gain of an oligomer upon tubulin binding (ΔG_olig_) was calculated from the concentrations of oligomers with different sizes (Extended Data Fig. 6j–l). The condition of chemical equilibrium reads

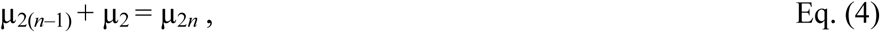

where

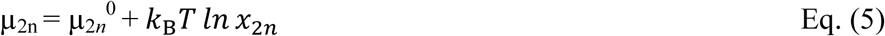

is the chemical potential of 2n-mer, with *µ*_2n_^0^ representing the concentration-independent part. By substituting Eq. (5) into Eq. (4), the free energy change upon the assembly of 2n-mer from 2(n-1)-mer can be represented by

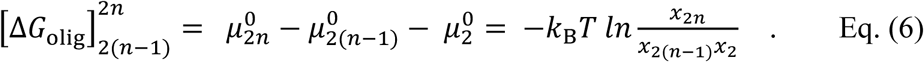

Based on the concentrations of oligomers with various sizes *x*_2_, *x*_4_, *x*_6_ … (Extended Data Fig. 6j–l), the values 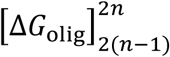 were obtained (Fig. 3i).

Exponential decay of *x*_2*n*_ for WT(GTP) and Y222F(GDP) tubulin indicates that with these tubulins, ΔG_olig_ is constant independent of the size of oligomer (2n). Thus, ΔG_olig_ was calculated from the slope of the lines in Extended Data Fig. 6j and l and plotted in Fig. 3i. The error bar indicates the error of the fit. For Y222F(GTP) tubulin, ΔG_olig_ for each 2n-mer was calculated by using Eq. (6), and the averages of the two experiments were plotted in Fig. 3i (no error bars).

In analogy with Eq. (6), when a MT containing 2(N-1) tubulin dimer incorporates another dimer, the free energy change upon binding should be obtained from the ratio 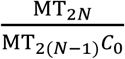 as

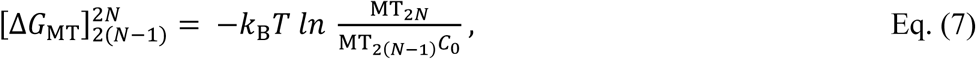

where C_0_ is the concentration of tubulin dimer, where we have ignored the consummation of dimer in oligomers. On the other hand, the binding equilibrium is determined by the rate constant for association and dissociation of tubulin dimer (*k*_+_ and *k*_-_) at the growing end of a MT,

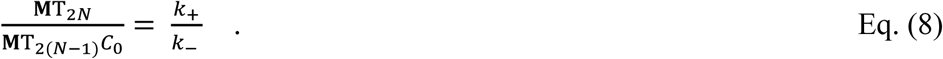

The rate constants, *k*_+_ and *k*_-_, can be calculated from the concentration dependence of the growth rate of MT, *v*_0_ = *k*_+_*C*_0_ − *k*_−_ (= number of tubulin dimers added onto each MT filament per sec), measured by in vitro MT dynamic assay (Extended Data Fig. 5e, g). Thus, the free energy gain upon binding of a tubulin dimer to a MT was calculated by the following equation and shown in Fig. 3i:

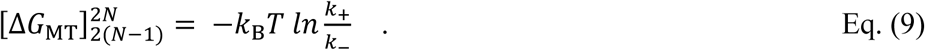

The error bars for each ΔG_MT_ in Fig. 3i represent the errors estimated from the standard deviation of 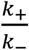. It is worth noting that while the energy changes 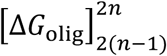 and 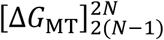 can contain an additive correction related to the choice of the standard volume for the mixing entropy calculation, the difference between these two energies, which should reflects the free energy change associated with the lateral interaction between tubulin dimers, does not suffer from it. The difference gives the absolute meaning.

### Statistical methods

Data analyses were conducted in Microsoft Excel 2013 (Microsoft office), Kaleidagraph 4.5 (Hulinks) and *ad hoc* programs written by Mathematica 8.0.4. (Wolfram Research).

### Reporting summary

Further information on research design is available in the Nature Research Reporting Summary.

### Data availability

The atomic coordinates and structure factors have been deposited in the Protein Data Bank under accession codes 6TIS (WT(GDP)), 6TIY (WT(GMPCPP)), 6TIZ (Y222F(GDP)) and 6TIU (Y222F(GTP)). The data that support the findings of this study are available from the corresponding authors upon reasonable request.

## Acknowledgements

This work is supported by the Grants-in-Aid for Scientific Research from the Ministry of Education, Culture, Sports, Science and Technology of Japan, 17H03668 (EM, SK, HI). A grant from the Foundation ARC pour la Recherche sur le Cancer (BG) is also acknowledged. We acknowledge the Chuo University Electron Microscopy Facility and the technical assistance by Ms. Nakahara (Chuo University) for the negative stain EM. Part of the cryo-EM study was conducted at the Advanced Characterization Nanotechnology Platform of the University of Tokyo, supported by the Nanotechnology Platform of the Ministry of Education, Culture, Sports, Science and Technology (MEXT), Japan. Diffraction data were collected at the SOLEIL synchrotron (PX1 and PX2 beam lines, Saint-Aubin, France) and the European Synchrotron Radiation Facility (ID23-1 beam line, Grenoble, France). We are most grateful to the machine and beam-line groups for making these experiments possible. Special thanks are also due to Dr. Masayuki Kajitani (Teikyo University) for providing tabacco mosaic viruses, the Drosophila Genomics Resource Center (supported by NIH grant 2P40OD010949) for providing *Drosophila* tubulin genes, Dr Yuichiro Maeda (Nagoya University) and Dr Takahide Kon (Osaka University) for critical reading of the manuscript, Dr. Hiroko Takazaki and Ms. Yoshimi Asano for assistance in sample preparation, Dr. Valérie Campanacci (I2BC) for image analysis and Dr. Marcel Knossow (I2BC) and the late Fumio Oosawa for their discussion and continuous encouragement.

## Author contributions

I.M., E.M. and B.G. designed the Y222F mutant and the experiments. S.U. and R.A. developed the experimental system for the expression and purification of *Drosophila* tubulin. S.I., H.I., R.A. and S.K. performed negative stain EM. S.I., H.I., M.S. and H.S. performed the cryo-EM work and S.I., R.A., K.X.N., M.H. and B.G. conducted oligomer image analysis. B.G. and T.M. conducted crystal structure analysis. R.A., E.M. and K.S. conducted turbidimetry and kinetic analysis, and M.H. measured the in vitro MT dynamics. E.M. and K.S. conducted statistical analyses of oligomers and built a model. E.M., K.S. and B.G. prepared the manuscript. All co-authors discussed the results and commented on the manuscript.

## Additional information

Correspondence and requests for materials should be addressed to Etsuko Muto, Benoît Gigant, Ken Sekimoto. Reprints and permissions information is available at www.nature.com/reprints.

---

**Extended Data Figure 1.**
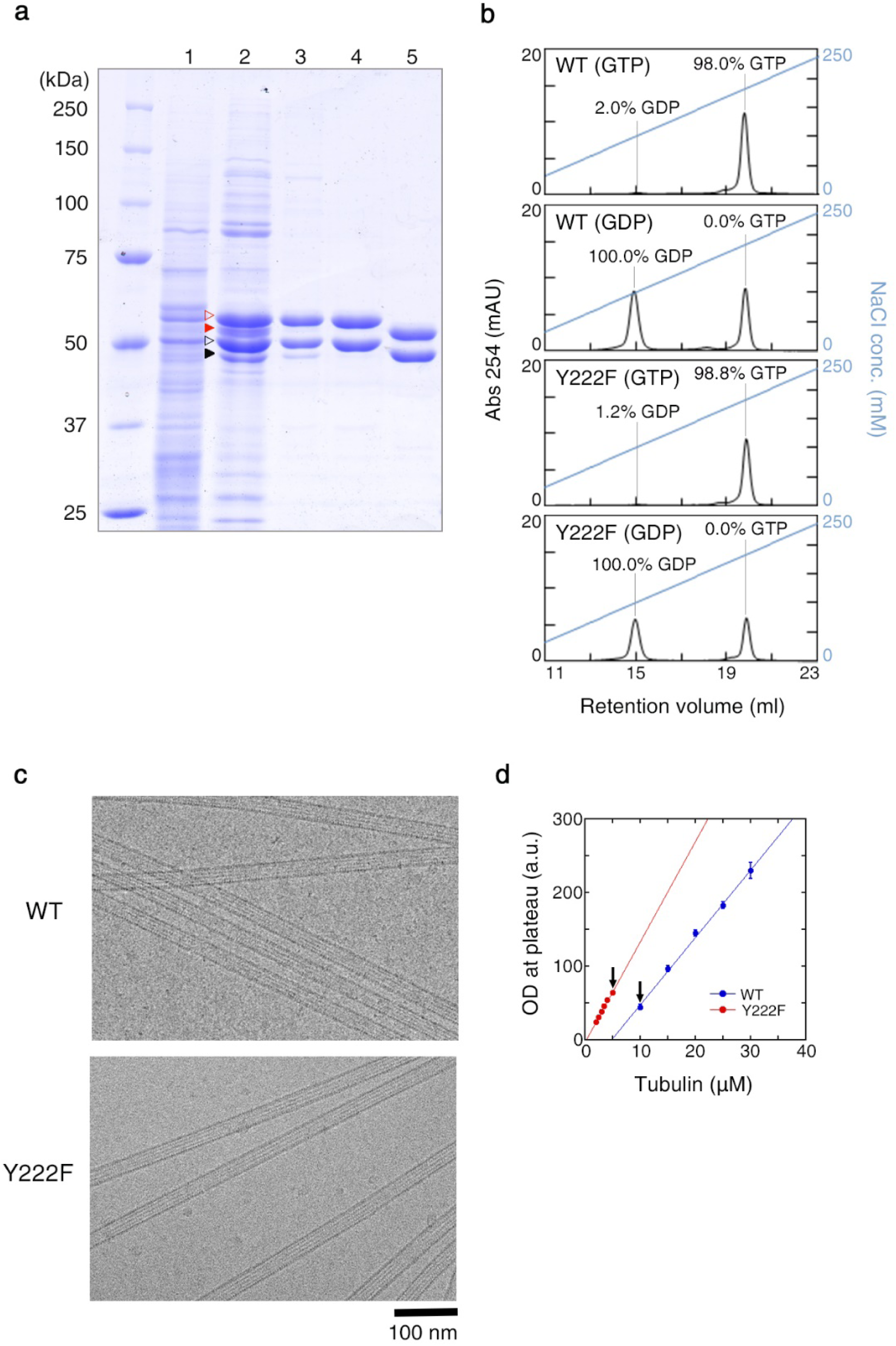
Preparation of *Drosophila* tubulin. **a**, SDS PAGE showing each step in preparation of WT tubulin. Lanes: (1) cell lysate, (2) DEAE anion exchange column eluent. **▷** and **▷**, recombinant α- and β-tubulin; ▶ and ▶, endogenous α- and β-tubulin, respectively. (3) His-tag affinity column eluent, (4) FLAG-tag affinity column eluent, (5) final product after tag cleavage. **b**, Analysis of nucleotide contents by FPLC. The percentage indicates the occupancies of the exchangeable site (E-site) calculated from each chromatogram, assuming equal numbers of exchangeable and nonexchangeable nucleotide sites per tubulin dimer and that the nonexchangeable nucleotide is GTP. mAU = milli-absorbance unit. **c**, Cryo-electron microscopy images of WT and Y222F MTs. **d**, Plateau value of turbidity (Fig. 1i, j) plotted as a function of the initial tubulin concentration, showing the critical concentration (= x-intercept) of WT and Y222F tubulin to be 4.7 ± 0.5 and 0.19 ± 0.15 (μM), respectively. The arrowheads indicate the pair of experiments compared in Fig. 2.

**Extended Data Figure 2.**
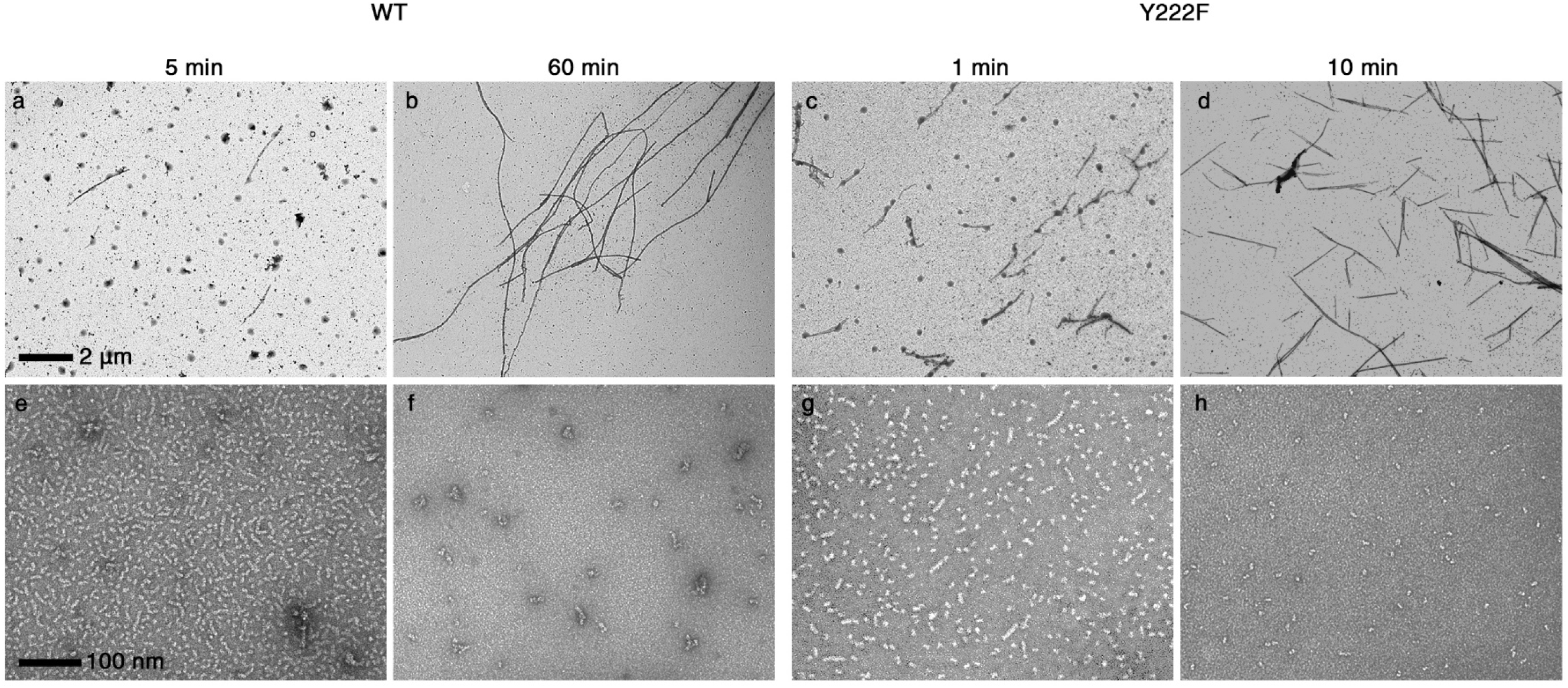
The number of oligomers decreased with the progress of MT assembly. Electron microscopy images of the WT MTs (**a, b**) and oligomers (**e, f**), Y222F MTs (**c, d**) and oligomers (**g, h**) recorded in the course of polymerization (Fig. 1i, j). The initial concentrations of tubulin were 10 μM (WT) and 5 μM (Y222F). The photographs of the MTs and oligomers were taken at 2,500x and 40,000x magnification, respectively. For Y222F mutant tubulin, only the dimers and aggregates are seen at 10 min, which is when the turbidity reached approximately 90% of the plateau value (h). In contrast, for WT tubulin, the oligomers and aggregates existed at 60 min, which is when the turbidity reached a plateau (f).

**Extended Data Figure 3.**
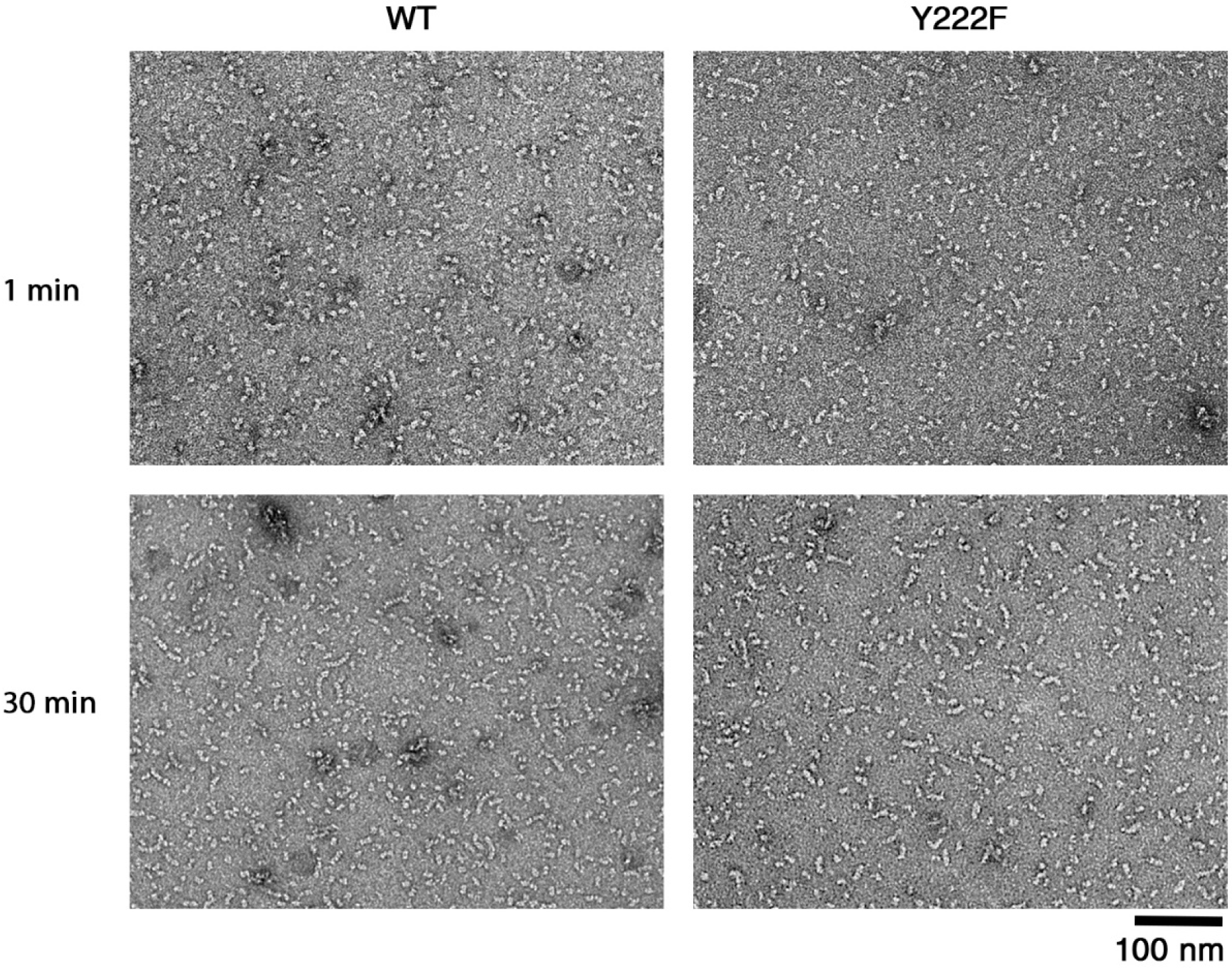
Electron microscopy images of the tubulin sample used for turbidimetry. Electron microscopy images of WT and Y222F tubulin 1 min after the ultracentrifugation (top) and 30 min later (bottom) at 4°C. Although our ultracentrifugation condition (see Methods) should sediment oligomers that are larger than dimers, we sometimes observed a tetramer in the Y222F sample immediately after ultracentrifugation, which may have been assembled during the 1 min needed for the preparation of the EM grid.

**Extended Data Figure 4.**
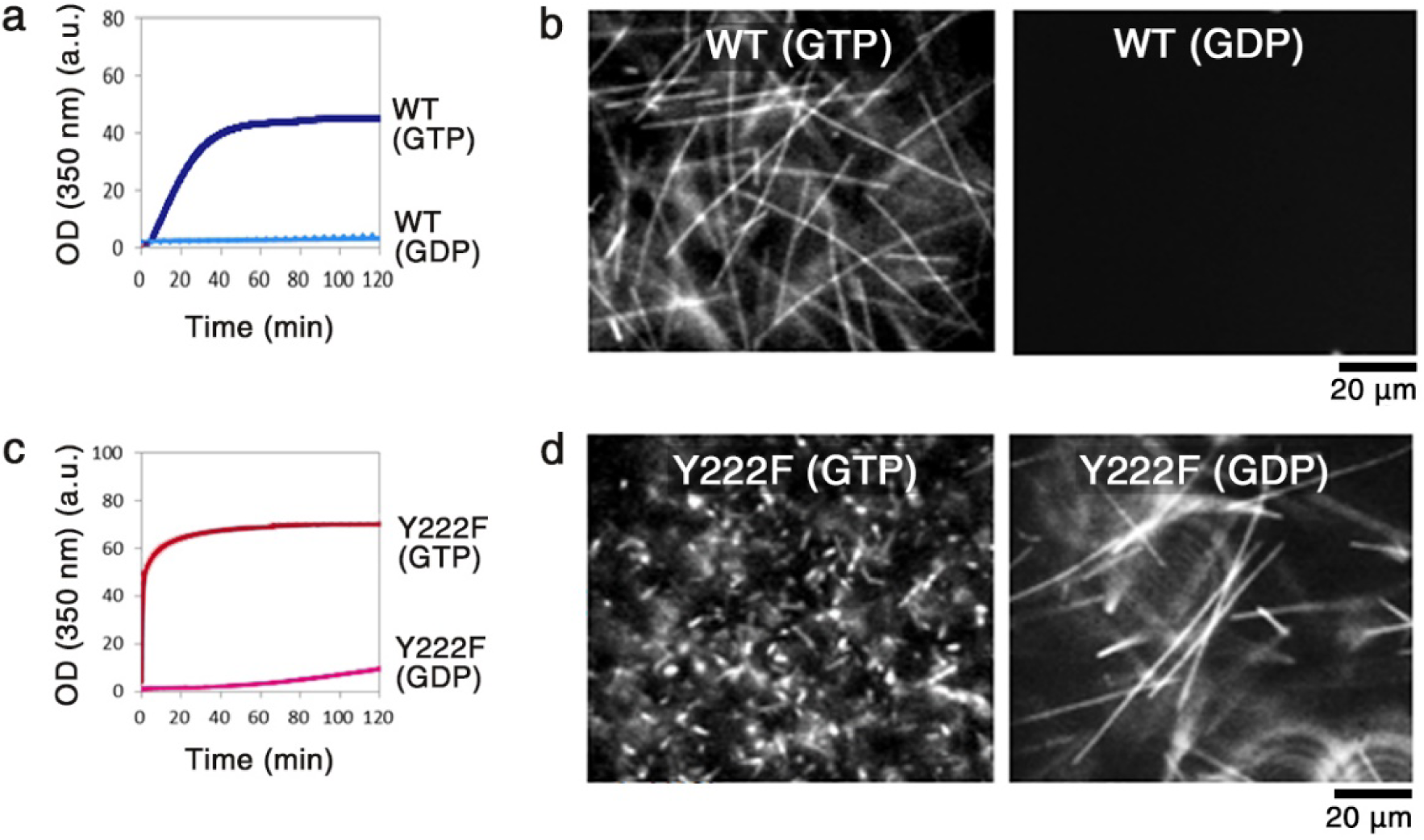
GDP-dependent assembly of Y222F tubulin. **a** and **c**, Time course of the polymerization of WT (10 μM) (a) and Y222F tubulin (6 μM) (c) monitored by OD_350_. **b** and **d**, Darkfield micrographs of the WT (b) and Y222F MTs (d) imaged at the end of reaction (120 min). While the WT(GTP) and Y222F(GTP) MTs were diluted for the optimal observation of individual MT filaments, Y222F(GDP) MTs were observed without dilution.

**Extended Data Fig. 5.**
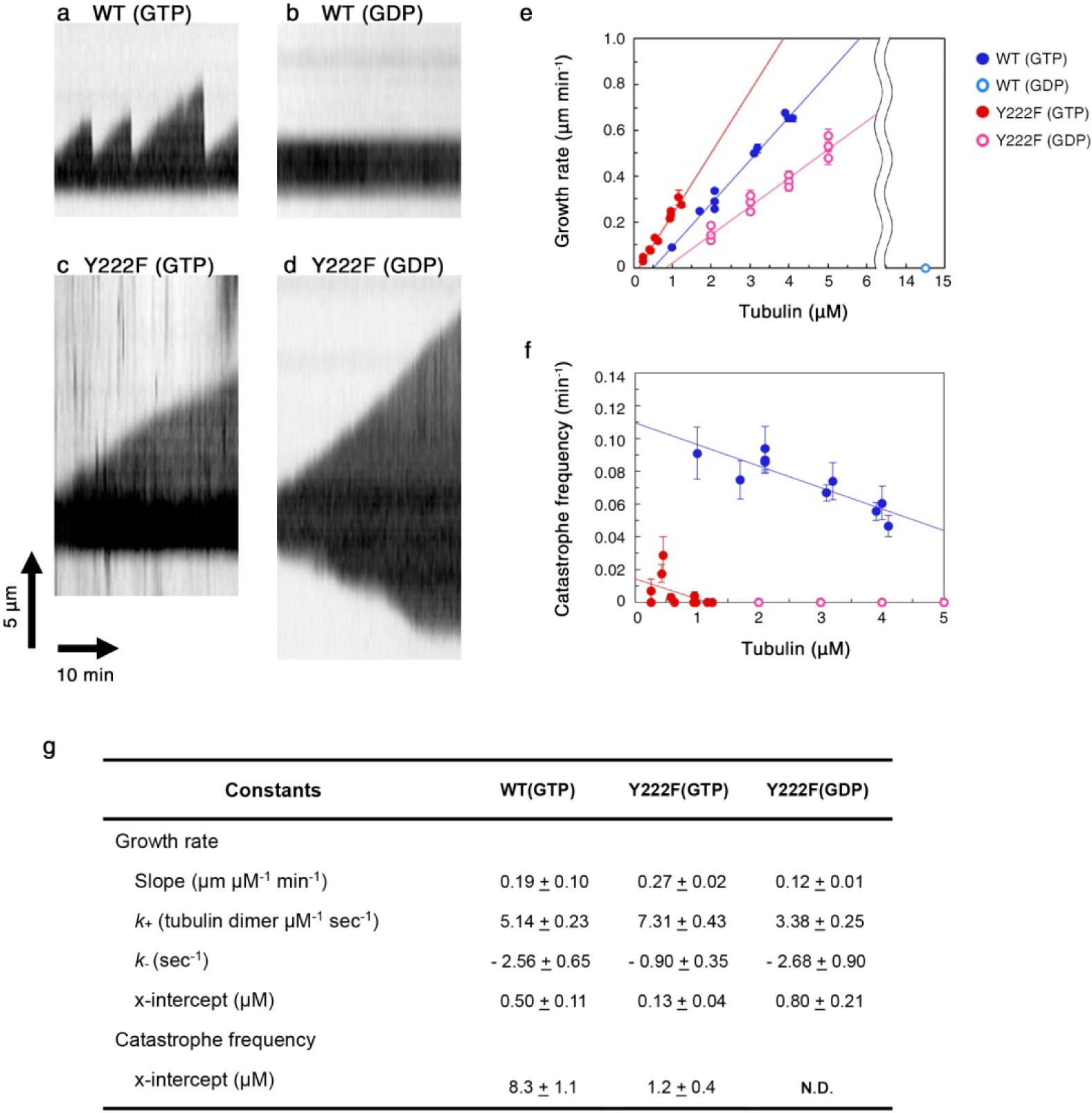
Dynamic instability of individual WT and Y222F MTs. **a** and **c**, Kymographs of WT and Y222F MTs polymerized in the GTP condition (tubulin concentration of 2.1 and 1.3 µM, respectively). **b** and **d**, Kymographs of WT and Y222F MTs polymerized in the GDP condition (8.4 and 3.0 µM, respectively). **e** and **f**, Concentration dependence of the growth rate (e) and frequency of catastrophe (f). In the case of Y222F(GDP), the growth rate and the catastrophe frequency at the plus end are reported. In panel (e), the regression line for WT(GTP), Y222F(GTP) and Y222F(GDP) can be represented by the equations, y =0.19 x – 0.09 (R = 0.99), y = 0.27x – 0.03 (R = 0.98), and y = 0.12x – 0.10 (R = 0.97), respectively. Total number of dataset was 966, 137, and 352 for WT(GTP), Y222F(GTP) and Y222F(GDP), respectively. In panel (f), the regression line for WT(GTP) and Y222F(GTP) can be represented by the equations y = –0.013 x + 0.109 (R= 0.88) and y = –0.012 x + 0.014 (R=0.46), respectively. Total time of elongation per data point was 112–2,491 min. **g**, Kinetic and thermodynamic parameters calculated from the data shown in (e) and (f). Values represent the mean ± s.d.

**Extended Data Fig. 6.**
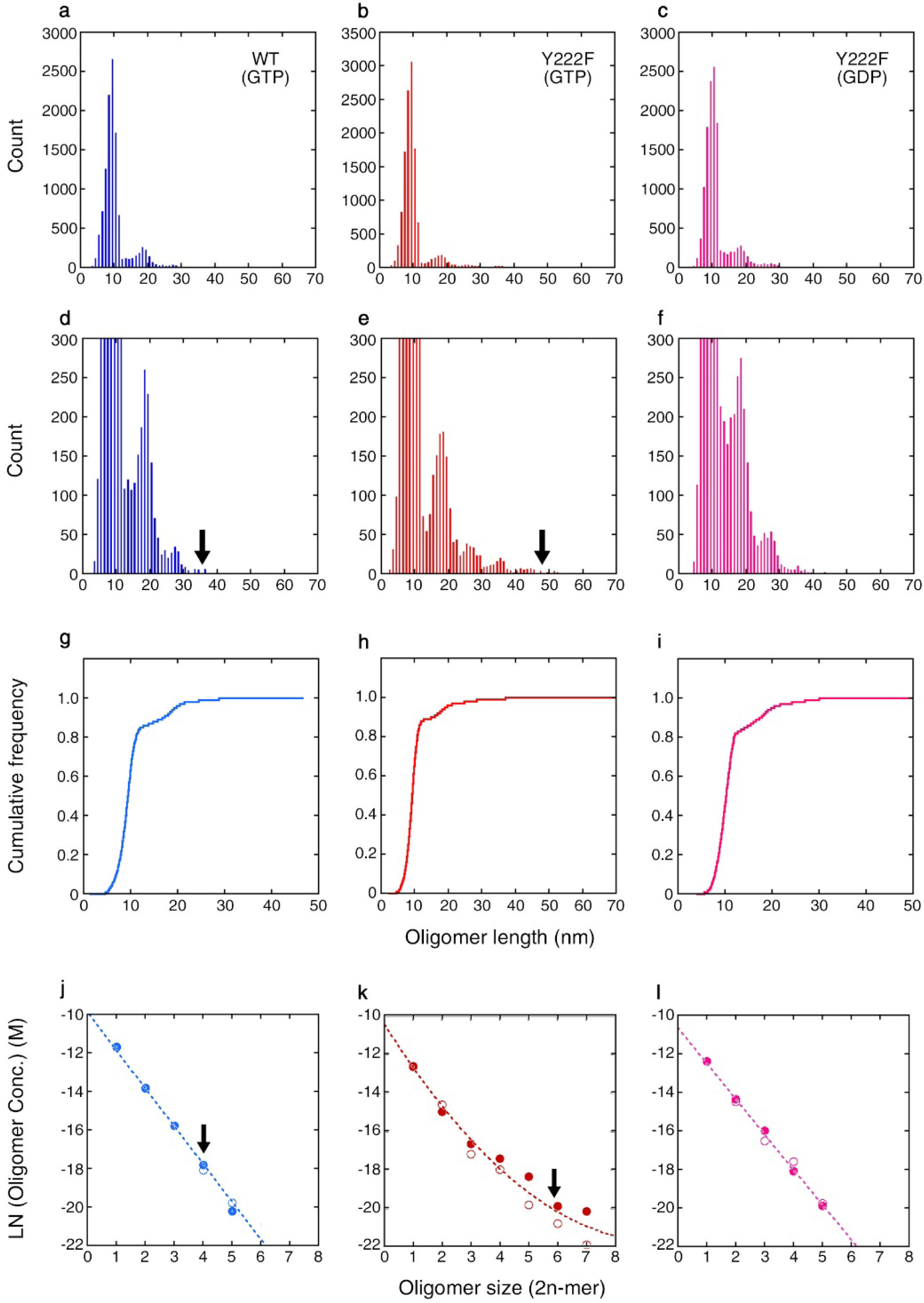
Size distribution of oligomers. **a–f,** The length distribution of WT(GTP) oligomers (a, d), Y222F(GTP) oligomers (b, e), and Y222F(GDP) oligomers (c, f). The same samples used for the curvature analysis (Fig. 2e) were analysed, but this time, the length of all oligomers, including dimers, was measured. In panels d–f, the scales for the vertical axes are enlarged. N = 11,527, 12,666, and 12,430 for the WT(GTP), Y222F(GTP), and Y222F(GDP) oligomers, respectively. **g–i**, Cumulative frequency of the length, calculated from the length distribution. **j–l**, Logarithm of the concentration of the oligomers, *x*_2*n*_, calculated from the probability of oligomers with different sizes (see Methods for the details of the calculation). The measurement was made twice for each type of tubulin (represented by filled and open circles). The raw data for the first round of measurement are shown in panels a**–**f. For the second round of measurement, N = 10,945, 11,803, and 11,092 for WT(GTP), Y222F(GTP) and Y222F(GDP), respectively. For both WT(GTP) and Y222F(GTP) tubulins, the largest oligomer was only one unit larger than the size of the critical nucleus, as estimated by the kinetic analyses in Fig. 4c. The size of the critical nucleus is indicated by arrows (panels d, e, j, k). The data for the WT(GTP) (j) and Y222F(GDP) oligomers (l) showed the exponential decay of the oligomer concentrations with size, which can be fit by the equations y = – 9.89 – 1.96 x (R = 0.99) and y = –10.63 – 1.83 x (R = 0.99), respectively. In the case of the Y222F(GTP) oligomers (k), the concentrations of oligomers with n>3 were significantly higher than the concentrations expected from simple exponential decay and were best fit by the equation y = –10.50 – 2.37 x + 0,13 x^2^ (R = 0.98).

**Experimental Data Fig. 7.**
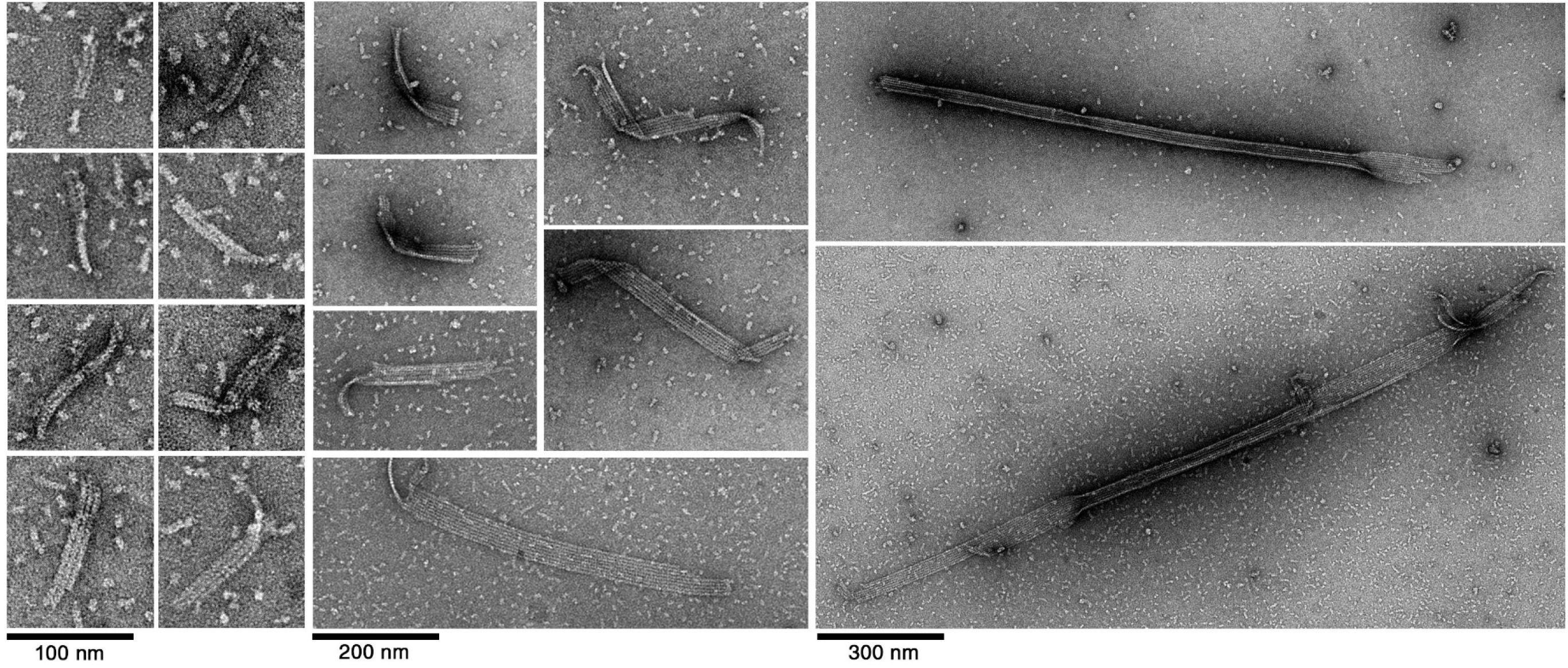
Electron microscopy images of the intermediate structures in the Y222F mutant. Multistranded oligomers and sheets observed at 30–120 sec after the onset of the reaction.

**Experimental Data Fig. 8.**
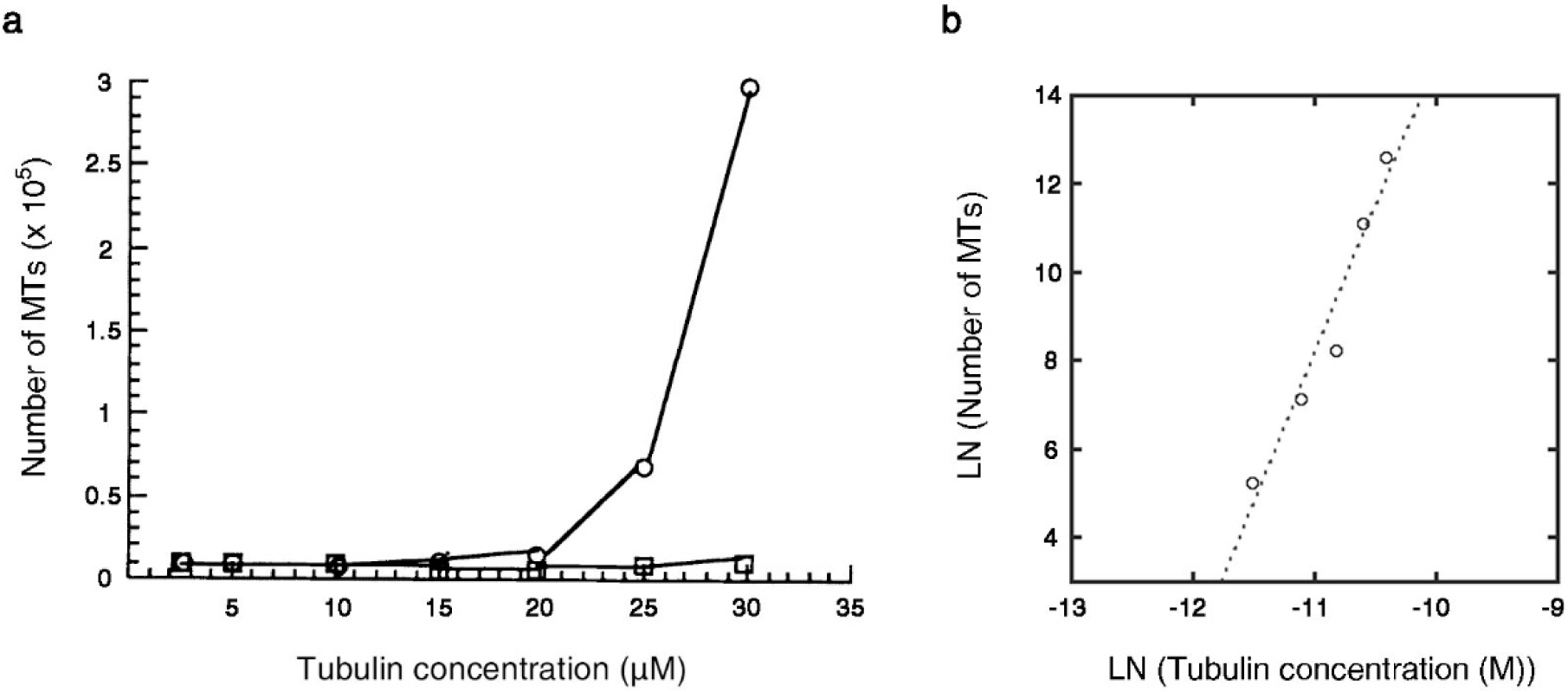
Initiation of MTs by the γTuRC template. **a**, Data from Fig. 3a in Zheng et al. (1995)^**14**^. The number of MTs newly formed within 5 min of incubation at 37°C with and without γTuRC (circle and square, respectively) at variable tubulin concentrations. **b**, Dataset for γTuRC-dependent nucleation plotted on a log-log scale. The data can be fit by the equation y = 81.5 + 6.7x (R = 0.97), indicating that the nucleation of MTs requires the formation of a critical nucleus composed of ∼7 tubulin dimers. We could not estimate the size of the critical nucleus for spontaneous nucleation (without γTuRC) because MTs were barely formed. [Credit: (a) Reprinted from Ref. 14 by permission from Springer Nature, Copyright (2020)].

**Extended Data Table 1.** Data collection and refinement statistics

|  | WT(GDP) | WT(GMPCPP) | Y222F(GDP) | Y222F(GTP) |
| --- | --- | --- | --- | --- |
| **Data collection\*** |  |  |  |  |
| Space group | P2<sub>1</sub>2<sub>1</sub>2<sub>1</sub> | P2<sub>1</sub>2<sub>1</sub>2<sub>1</sub> | P2<sub>1</sub>2<sub>1</sub>2<sub>1</sub> | P2<sub>1</sub>2<sub>1</sub>2<sub>1</sub> |
| Cell dimensions<br>*a*, *b*, *c* (Å) | 66.6 126.5 250.2 | 66.7 126.6 251.0 | 66.2 126.7 250.4 | 66.0 126.3 249.8 |
| Resolution (Å) | 48 - 2.30<br>(2.34-2.30) | 48.2-2.29<br>(2.33-2.29) | 63.4 – 2.20<br>(2.23-2.20) | 50.0 – 3.57<br>(3.66-3.57) |
| R<sub>meas</sub> | 0.092 (1.13) | 0.112(1.87) | 0.125 (1.25) | 0.51 (3.54) |
| *I*/σ*I* | 10.3 (1.0) | 13.2 (1.2) | 12.8 (1.6) | 4.1 (0.7) |
| CC 1/2 | 0.999 (0.611) | 0.999 (0.612) | 0.998 (0.912) | 0.979 (0.41) |
| Completeness (%) | 98.8 (92.1) | 99.5 (89.8) | 99.5 (91.2) | 98.7 (83.1) |
| Multiplicity | 4.1 (3.4) | 7.4 (6.6) | 9.4 (6.2) | 11.7 (8.9) |
| **Refinement** |  |  |  |  |
| Resolution (Å) | 2.30 | 2.29 | 2.20 | 3.57 |
| No. reflections | 87,343 | 91,616 | 104,968 | 25301 |
| *R*<sub>work</sub>/*R*<sub>free</sub> | 0.183/0.218 | 0.187/0.217 | 0.167/0.208 | 0.221/0.244 |
| No. atoms |  |  |  |  |
| Protein | 14509 | 14560 | 14453 | 14483 |
| Ligand/ion | 189 | 193 | 199 | 185 |
| Water | 550 | 432 | 759 | 0 |
| B-factors |  |  |  |  |
| Protein | 68.1 | 64.2 | 52.6 | 111 |
| Ligand/ion | 75.6 | 75.0 | 68.6 | 88.7 |
| Water | 63.7 | 60.1 | 54.0 |  |
| R.m.s deviations |  |  |  |  |
| Bond lengths (Å) | 0.008 | 0.008 | 0.010 | 0.008 |
| Bond angles (°) | 0.93 | 0.98 | 1.08 | 0.98 |
| Coordinate error (Å) | 0.33 | 0.32 | 0.26 | 0.60 |
| Ramachandran (%) |  |  |  |  |
| Favored region | 97.48 | 97.50 | 96.87 | 96.77 |
| Allowed region | 2.02 | 2.12 | 2.91 | 2.96 |
| Outliers | 0.49 | 0.38 | 0.22 | 0.27 |
| **PDB Code** | 6TIS | 6TIY | 6TIZ | 6TIU |

\*Data were collected on a single crystal. Highest resolution shell is shown in parenthesis.

